# Single cell-resolved transcriptional dynamics of human subcutaneous adipose tissue during lifestyle- and bariatric surgery-induced weight loss

**DOI:** 10.1101/2025.01.30.634294

**Authors:** Anne Loft, Rasmus Rydbirk, Ellen Gammelmark Klinggaard, Elvira Laila Van Hauwaert, Charlotte Wilhelmina Wernberg, Andreas Fønss Møller, Trine Vestergaard Dam, Mohamed Nabil Hassan, Babukrishna Maniyadath, Ronni Nielsen, Aleksander Krag, Joanna Kalucka, Søren Fisker Schmidt, Mette Enok Munk Lauridsen, Jesper Grud Skat Madsen, Susanne Mandrup

## Abstract

During sustained weight gain, human white adipose tissue undergoes dramatic remodeling that may compromise adipose tissue function and lead to obesity comorbidities such as cardiometabolic disease. These comorbidities can be at least partially reversed by weight loss; however, the degree to which human adipose tissue is remodeled during weight loss is currently unclear. Here, we have used snRNA-seq combined with bulk RNA-seq and 3D light microscopy to investigate the cellular and transcriptional alterations in abdominal subcutaneous adipose tissue (ASAT) from a unique cohort of obese individuals undergoing initially modest (8-10%) lifestyle-induced weight loss followed by a more dramatic (20-45%) bariatric surgery-induced weight loss. We show that in response to this weight loss, ASAT in both males and females underwent dramatic compositional and transcriptional remodeling. Most notably, surgery-indued weight loss led to an increase in the adipose vascular compartment, a dramatic reduction in proinflammatory immune cells, and a tissue-wide reduction in inflammatory gene signatures. In response to modest weight loss, we observed an increase in specific progenitor populations and an induction of proadipogenic genes, indicating that this process is important for the early adaptation of AT to weight loss that might contribute to the early overall beneficial effects on insulin sensitivity and systemic inflammation.

## Introduction

White adipose tissue (WAT) is a highly plastic tissue that plays a major role in systemic energy homeostasis by serving as the major depot for storage of metabolic energy as well as the endocrine source of metabolic hormones ^1^. The critical importance of well-functioning adipocytes is clearly demonstrated by the severe metabolic comorbidities associated with lipodystrophies commonly characterized by the inability to properly store lipids in adipose tissues (AT) ^2^. On the other hand, excessive accumulation of AT, i.e., obesity, is also associated with an increase in AT dysfunction and systemic comorbidities, especially when the expansion is driven by enlargement of the existing adipocytes (hypertrophy) rather than *de novo* formation of new fat cells (hyperplasia). A large body of literature has shown that adipose hypertrophy is associated with substantial changes in the cellular composition and architecture of AT, including recruitment and activation of pro-inflammatory myeloid cells as well as remodeling of the extracellular matrix and the vascular cell compartment ^1,3^. Studies based on single cell RNA sequencing (scRNA-seq) and single nuclei RNA sequencing (snRNA-seq) have recently provided further insight into the cellular composition of AT and the plasticity during weight gain in mouse models as well as humans ^4–8^. These studies have revealed that obesity is associated with major changes in cellular subtypes, especially in the immune cell compartment, where there is a pronounced increase in the lipid-associated macrophages (LAMs). Moreover, obesity is associated with major changes in gene expression in all cell types, including a general increase in proinflammatory signals in multiple cell types ^4,7^.

Weight loss has been reported to reverse many the cardiometabolic comorbidities at least partially ^9^; however, there is considerable disagreement regarding the extent to which AT inflammation is reversed. Studies in mice using calorie restriction or low-fat diet to reverse high-fat diet-induced obesity have indicated an initial increase followed by a decrease in macrophage infiltration in AT^10,11^. In humans, short-term lifestyle-induced weight loss has been reported to either decrease ^12^, increase ^13,14^, or have no effects ^15–17^ on AT inflammation; however, overall, it seems that the beneficial systemic changes, such as increased insulin sensitivity and decrease in systemic inflammation, precedes reversal of AT inflammation ^15,18,19^. While short-term effects of weight loss on AT inflammation differ between studies, long-term major lifestyle-induced weight loss has almost invariably been associated with a decrease in AT inflammation in humans ^20,21^. Similarly, the major weight loss induced by bariatric surgery has been reported to decrease inflammation in human subcutaneous AT and increase the number of adipogenic precursor cells that might support a healthy tissue remodeling following a pronounced weight loss ^22–24^. Recently, snRNA-seq was applied to study cellular remodeling in omental and subcutaneous AT in individuals (2 males, 3 females) undergoing bariatric surgery ^25^. Notably, this study did not identify consistent cellular compositional differences between AT isolated during surgery and AT isolated 2 years post-surgery, indicating that, despite the reduction in AT inflammation reported in other studies, compositional changes in AT may be relatively modest.

Here, we have used snRNA-seq combined with bulk-seq and bioimaging to investigate the cellular and transcriptional alterations in abdominal subcutaneous AT (ASAT) from a unique cohort of obese individuals that underwent initially modest (8-10%) lifestyle-induced weight loss followed by a more dramatic (20-45%) bariatric surgery-induced weight loss. We show that in response to weight loss, in particular the surgery-induced weight loss, ASAT in both males and females underwent dramatic compositional, structural and transcriptional remodeling including decreased inflammation and an apparent increase in angiogenic and adipogenic potential.

## Results

### Overall cell type-resolved map of the human adipose tissue in response to lifestyle- and surgery-induced weight loss

To investigate the cellular and molecular transitions in the AT in response to lifestyle- and surgery-induced weight loss, we recruited obese patients (BMI>35) enrolled for bariatric surgery to a longitudinal clinical study (the ATLAS study) (Fig. 1a) (see Methods for inclusion and exclusion criteria). ASAT biopsies were sampled at the time of enrollment (visit 1), the time of bariatric surgery following a mandatory lifestyle-induced weight loss (visit 2), and two-years post-surgery (visit 3) (Fig. 1a). All patients underwent detailed clinical characterization (Table 1), from which we selected 7 females and 7 males with the highest metabolic syndrome (MetS) scores at visit 1 for single cell sequencing analyses of ASAT biopsies. A significant (5-10%) lifestyle-induced weight loss was observed at the time of bariatric surgery, and this was followed by a bariatric surgery-induced weight loss that increased gradually over the first year following surgery (20-45%) (Fig. 1b-c). The weight loss was comparable in both sexes and remained stable until the third biopsy was collected two years post-surgery (Fig. 1c). Both sexes experienced a gradual decline in BMI and body fat percentage in response to the progressive weight loss, alongside an improvement in HOMA-IR and some blood lipid parameters following bariatric surgery (Fig. 1d-f and Table 1).

**Fig. 1:**
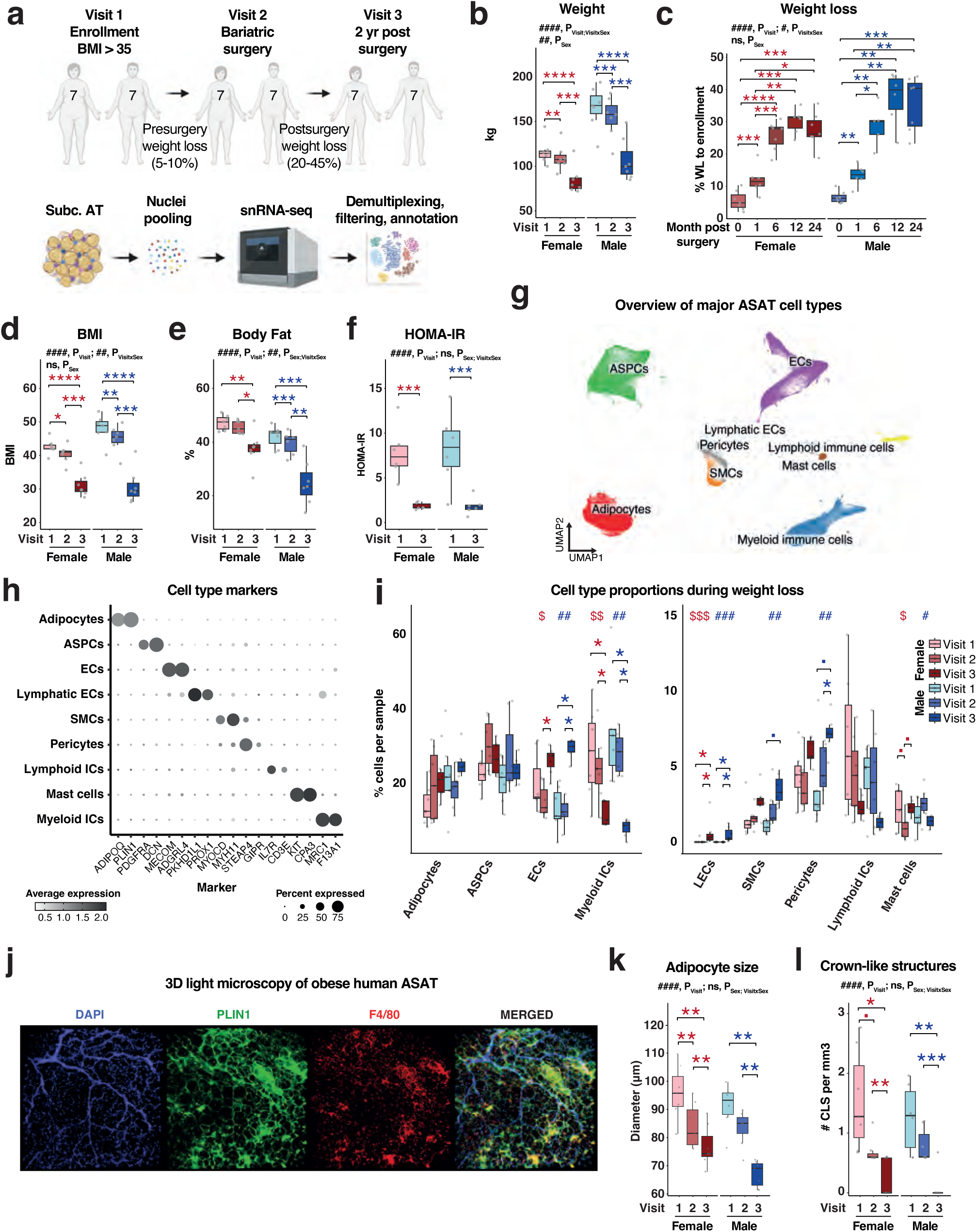
a, ASAT biopsies were collected from 7 females and 7 males with severe obesity (BMI>35) at the time of enrollment (visit 1), during bariatric surgery (visit 2) and 2 yr post-surgery (visit 3). Nuclei isolated from ASAT biopsies from different donors were pooled, single cell-sequenced using the 10X genomics platform and genetically demultiplexed and filtered to only retain high quality nuclei. b-f, Weight (b), weight loss (c), BMI (d), body fat (e), and HOMA-IR (f) at the indicated visits (female, n=6-7, male, n=6-7). g, Uniform manifold approximation and projection (UMAP) of 130,918 ASAT nuclei from all donors at all 3 visits. h, Marker genes for each cell population in the human ASAT dataset. i, The proportion of a specific cell population relative to all cells in the human ASAT dataset. j, representative images obtained using 3D light sheet immunofluorescence microscopy of DAPI (blue), PLIN1 (green), F4/80 (red) and merged image at visit 1. k, quantification of the average adipocyte diameter in ASAT (female, n=7; male, n=6-7). l, quantification of the number of crown-like structures (CLS) per mm^3^ tissue (female, n=7; male, n=6-7). Box plot data: center line, median; box limits, upper and lower quartiles; whiskers, 1.5x IQR. A repeated measures 2-way Anova (in b, d, and f) or mixed-effects analysis (in c, e, k and l), followed by post hoc Tukey’s multiple comparison test (in b-e, k-l) or uncorrected Fisher’s LSD test (in f) were used to evaluate the effects of visits and sex and their interaction on different clinical parameters. A Kruskal-Wallis (KW) test, followed by a post-hoc Wilcoxon signed-rank test with Holm’s correction for multiple testing, was performed to assess significance between visits for each sex separately in i (▪, p<0.1; #/$/*, p<0.05; ##/$$/**, p<0.01; ###/$$$/***, p<0.001; ####/****, p<0.0001). ASPCs, adipogenic stromal and progenitor cells; ECs, endothelial cells; ICs, Immune Cells; SMCs, (vascular) smooth muscle cells. Credit: a, Created with BioRender.com.

**Table 1.**

|  | Females |  |  |  | Males |  |  |  |
| --- | --- | --- | --- | --- | --- | --- | --- | --- |
|  | Visit 1 (N=7) | Visit 2 (N=7) | Visit 3 (N=7) | P-value | Visit 1 (N=7) | Visit 2 (N=7) | Visit 3 (N=7) | P-value |
| **Age [y]** | 53.4 ± 3.4 | 53.9 ± 3.6 | 56.4 ± 3.9 | - | 41.3 ± 13.0 | 42.0 ± 12.8 | 43.9 ± 12.9 | - |
| **Time since enrollment [d]** | 0.0 ± 0.0 | 295.7 ± 92.8 | 1071.3 ± 89.2 | - | 0.0 ± 0.0 | 264.3 ± 79.7 | 977.7 ± 85.9 | - |
| **Body Mass Index (BMI)** | 42.5 ± 2.2 | 40.4 ± 2.6 | 31.2 ± 3.5 | #### | 48.0 ± 4.1 | 45.0 ± 3.9 | 30.9 ± 5.1 | #### |
| **Weight [kg]** | 116.0 ± 13.8 | 109.7 ± 14.3 | 84.6 ± 15.5 | § | 165.5 ± 23.7 | 154.7 ± 22.7 | 106.9 ± 23.8 | ### |
| **Waist [cm]** | 124.8 ± 9.3 | 114.5 ± 9.5 | 102.4 ± 11.7 | ## | 148.9 ± 12.1 | 137.3 ± 15.0 | 110.7 ± 13.6 | ### |
| **Waist-hip-ratio (WHR)** | 0.9 ± 0.0 | 0.9 ± 0.1 | 0.9 ± 0.0 | ns | 1.1 ± 0.0 | 1.0 ± 0.1 | 1.0 ± 0.1 | ns |
| **Body fat [%]** | 47.2 ± 2.8 | 45.5 ± 2.4 | 37.6 ± 6.3 | §§ | 42.0 ± 4.0 | 39.6 ± 4.4 | 24.8 ± 8.3 | #### |
| **Systolic BP [mmHg]** | 133.6 ± 13.2 | 120.2 ± 5.9 | 127.1 ± 16.0 | ns | 134.3 ± 16.3 | 140.2 ± 14.0 | 131.9 ± 16.0 | ns |
| **Diastolic BP [mmHg]** | 88.9 ± 9.9 | 82.5 ± 5.9 | 84.9 ± 11.1 | ns | 88.4 ± 5.8 | 90.2 ± 5.3 | 78.9 ± 13.6 | ns |
| **Blood glucose [mmol/L]** | 7.0 ± 1.2 | na | 5.8 ± 0.8 | \$ | 6.2 ± 0.6 | na | 5.1 ± 0.5 | \$ |
| **Insulin [U/mL]** | 161.0 ± 24.2 | na | 68.6 ± 49.8 | \$\$\$ | 214.5 ± 117.0 | na | 49.7 ± 24.3 | \$ |
| **HOMA IR** | 7.8 ± 2.9 | na | 1.8 ± 0.4 | \$\$ | 8.2 ± 4.1 | na | 1.8 ± 1.0 | \$\$ |
| **HbA1c [mmol/mol]** | 50.1 ± 12.1 | na | 40.4 ± 9.7 | ns | 37.0 ± 3.5 | na | 33.5 ± 3.6 | \$ |
| **Triglycerides [mmol/L]** | 1.8 ± 0.7 | na | 1.5 ± 0.8 | ns | 1.4 ± 0.3 | na | 1.0 ± 0.3 | \$ |
| **Total cholesterol [mmol/L]** | 4.1 ± 1.3 | na | 4.4 ± 0.8 | ns | 4.0 ± 1.1 | na | 3.9 ± 0.4 | ns |
| **HDL [mmol/L]** | 1.0 ± 0.2 | na | 1.5 ± 0.3 | \$\$ | 0.9 ± 0.1 | na | 1.4 ± 0.2 | \$\$ |
| **LDL [mmol/L]** | 2.6 ± 1.3 | na | 2.5 ± 0.8 | ns | 2.7 ± 1.2 | na | 2.3 ± 0.7 | ns |
| **MetS score** | 3.9 ± 1.1 | na | 3.0 ± 1.8 | ns | 3.4 ± 0.8 | na | 1.0 ± 0.9 | \$\$\$ |

Data shown is as mean±SD.
If data were available for all visits, a one-way paired ANOVA (#) or a Kruskal-Wallis (§) test were conducted for each sex individually.
If only data for visit 1 and 3 were available, a paired t-test (\$) or a Wilcoxon Signed Rank test (\*) were performed for each sex individually.

To mitigate potential inter-individual batch effects in the snRNA-seq analyses, we pooled nuclei isolated from AT from the 14 donors at each visit prior to single cell analyses (Fig. 1a). After genetic demultiplexing and extensive quality filtering, we recovered more than 130.000 nuclei and observed a similar number of detected unique molecular identifiers (UMIs) and genes across all samples (Extended Data Fig. 1a). Using *Conos* ^26^, we constructed a joint graph that allowed us to identify 9 distinct cell clusters and then applied uniform manifold approximation and projection (UMAP) to map these clusters into a two-dimensional space. The identified clusters were annotated as adipocytes, adipogenic stromal and progenitor cells (ASPCs), endothelial cells (ECs), lymphatic ECs, smooth muscle cells (SMCs), pericytes, lymphoid immune cells, mast cells, and myeloid immune cells based on marker gene expression profiles (Fig. 1g-h, Extended Data Table 1-2). The major cell populations in our dataset correspond to the populations identified in our previously published integrated sxRNA-seq atlas of human WAT (WATLAS ^8^) (Extended Data Fig. 1b). Nuclei from all visits, sexes, and donors are represented across all major clusters (Extended Data Fig. 1c-e). We next explored how the relative cell type composition of ASAT is affected by weight loss induced by lifestyle intervention and by bariatric surgery. The most dramatic changes in cell fractions occurred within the myeloid compartment, where bariatric surgery-induced weight loss caused a reduction in myeloid immune cells in both sexes, dropping from approximately 30% in obese individuals to less than 10% following surgery (Fig. 1i). By contrast, the moderate lifestyle-induced weight loss appeared to be insufficient to efficiently induce reversal of myeloid immune cell infiltration (Fig. 1i). We also observed remarkable fractional changes in the vascular compartment, where bariatric surgery-induced weight loss led to increases in both pericytes, SMCs and ECs (Fig. 1i). We did not observe any significant changes in the fraction of adipocytes (Fig. 1i), consistent with the notion that weight loss is primarily driven by a decrease in adipocyte volume^27–29^.

To investigate adipocyte cell size and validate the dramatic compositional changes in the myeloid compartments, we performed 3D light sheet immunofluorescence microscopy of adipose biopsies using antibodies for the adipocyte marker, PLIN1, and the myeloid marker, F4/80 (Fig. 1j, Extended Video 1-2). The PLIN1 staining allowed us to determine adipocyte size by 3D image analysis (>1000 cells per sample) (Fig. 1j, Extended Video 2) revealing a significant decrease in adipocyte diameter in ASAT already following lifestyle-induced weight loss and an even greater decrease in diameter following surgery-induced weight loss (Fig. 1k, Extended Data Fig. 1f). The number of crown-like structures (CLS), which represent dying adipocytes surrounded by macrophages, was not significantly reduced following lifestyle-induced weight loss, although a trend is noted. However, these structures are almost completely absent after surgery-induced weight loss, consistent with the low number of myeloid cells in the snRNA-seq data (Fig. 1l).

In summary, in this cohort of severely obese men and women, moderate lifestyle-induced weight loss over a period of 6 to 12 months did not lead to consistent changes in the overall cellular composition of ASAT. However, bariatric surgery and the ensuing dramatic weight loss led to a marked proportional decrease in myeloid cells as well as an increase in different types of vascular cells.

### Capillary and venular endothelial cells are enriched in adipose tissue after bariatric surgery

Vascularization of AT is crucial for tissue function ^30^. Recent single-cell studies have examined the effects of obesity on the endothelium in both mice and humans, revealing highly subpopulation-specific changes ^5,31,32^; however, the impact of weight loss on vascular subpopulations in human AT remains unexplored. The vascular compartment contains vascular and lymphatic ECs, SMCs, and pericytes marked by *MECOM, PROX1*, *MYH11,* and *STEAP4,* respectively (Fig. 1g-h). Further subclustering separated vascular ECs into arterial ECs (arECs), venous ECs (vEC), and capillary ECs (cECs) (Fig. 2a and Extended Data Fig. 2a-c). arECs are marked by *PCSK5,* and *NEBL* and vECs express high levels of *ACKR1* and *IL1R1* (Fig. 2b, Extended Data Table 1-2), displaying a strong association with previously annotated arECs and vECs, respectively ^8^ (Extended Data Fig. 2d). cECs are marked by *BTNL9* and *CADM2* and can be further distinguished into two subtypes, cEC and capillary venous ECs (cvECs), where cvECs additionally express low levels of venous markers and are strongly associated with both previously annotated capillary and venous ECs (Fig. 2b and Extended Data Fig. 2d). In line with the overall fractional increase in vascular ECs, the proportions of vECs in both sexes and of cECs in males were increased after bariatric surgery (Fig. 2c). Furthermore, velocity analyses suggest that transitions between vascular cells became much more pronounced following surgery-induced weight loss consistent with the notion that the sustained weight loss induced comprehensive vascular remodeling (Fig. 2d).

**Fig. 2:**
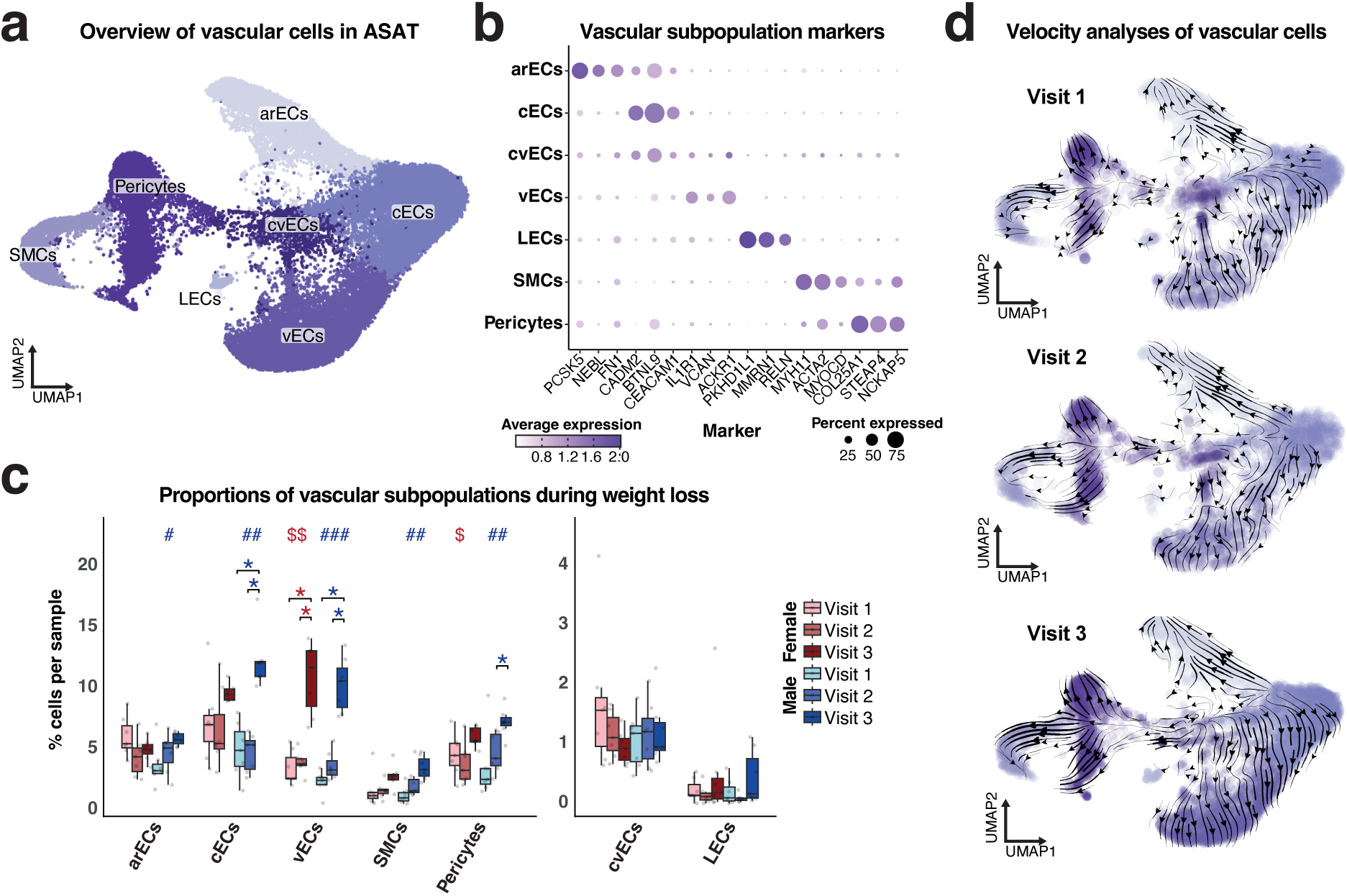
a, UMAP projection of vascular cells from all donors at all 3 visits. b, Marker genes for each vascular subpopulation. c, The proportion of a specific vascular subpopulation relative to all cells in the human ASAT dataset. d, UMAP plots visualizing RNA velocity of vascular cells at visit 1, 2 and 3. The arrows indicate the predicted future cell state. Box plot data: center line, median; box limits, upper and lower quartiles; whiskers, 1.5x IQR. A KW test, followed by post-hoc Wilcoxon signed-rank test with Holm’s correction for multiple testing, was performed to assess significance between visits for each sex separately in c (#/$/*, p<0.05; ##/$$, p<0.01; ###, p<0.001). arECs, arterial ECs; cECs, capillary ECs; cvECs, capillary venous ECs; LECs, lymphatic EndoCs; vECs, venous ECs; SMCs, (vascular) smooth muscle cells.

In summary, our results indicate that surgery-induced weight loss led to increased vascularization of ASAT as well as remodeling of the vascular compartment, persisting at least two years post-surgery.

### Weight loss induces extensive alterations in the immune cell compartment

Obesity is known to increase the presence of immune cells in AT, in particular macrophages, and the resulting low-grade inflammation is believed to contribute to AT dysfunction and systemic insulin resistance ^33–35^. Interestingly, however, the degree to which immune cells disappear or are transcriptionally remodeled by weight loss is not well understood. The immune cell compartment in ASAT from our human cohort comprises lymphoid immune cells (characterized by *CD3E* and *IL7R*), mast cells (characterized by *KIT* and *CPA3*), and myeloid immune cells (characterized by *MRC1* and *F13A1*), most of which exhibited a fractional decrease during surgery-induced weight loss (Fig. 1g-i). In line with this, expression of general markers of inflammation, such as *TNF*, *CCL5*, and *CCL3*, was significantly decreased in bulk RNA-seq data of ASAT from an extended patient cohort 2 years after bariatric surgery (Extended Data Fig. 3a). To investigate how different subpopulations of lymphoid and myeloid immune cells were changed after weight loss, we subclustered each of these cell types separately. Within the lymphoid compartment, we identified T-regs, CD4 and CD8 T-cells, NK-like cells, and NK cells (Extended Data Fig. 3b-f, Extended Data Table 1-2). Similar to the overall lymphoid immune cell population, the proportion of most of the lymphoid subpopulations in AT tended to decrease after bariatric surgery, although not reaching significance (Extended Data Fig. 3g). Within the myeloid immune cells, we identified several different subpopulations of monocytes, macrophages, and DCs (Fig. 3a and Extended Data Fig. 3h-j). The largest myeloid immune cell cluster, likely representing AT-resident (perivascular) macrophages (ATMs), was characterized by high *LYVE1* expression and was enriched for transcripts related to receptor-mediated endocytosis (Fig. 3b, Extended Data Fig. 3k, Extended Data Table 1-2). Furthermore, we annotated two types of lipid-associated macrophages (LAMs) that were both enriched for transcripts related to lipid- and cholesterol-handling pathways. The ‘classical’ LAM expressed high levels of canonical LAM-markers, such as *TREM2*, *LPL, FABP5,* and *SPP1* corresponding to previously reported LAM subpopulation in human AT (Fig. 3b and Extended Data Fig. 3k) ^8,36,37^. In contrast, early LAMs had lower expression of canonical LAM markers, but similar high expression of genes related to cholesterol metabolism, *CYP27A1* and *APOE (*Fig. 3b, Extended Data Table 1-2). We also detected smaller clusters of different subtypes of dendritic cells as well as a population of cells sharing markers of monocytes and macrophages denote mono/mac cells. For most subpopulations of myeloid cells, we found that bariatric surgery, but not moderate weight loss, led to a proportional reduction (Fig. 3c), suggesting that a prolonged and dramatic weight loss is required to remodel the myeloid compartment in AT. The LAM subpopulations were particularly depleted after bariatric surgery, and consistent with this, the LAM marker genes, *TREM2*, *SPP1*, and *MMP9* were expressed at very low levels in ASAT from an extended patient cohort 2 years after surgery (Fig. 3d).

**Fig. 3:**
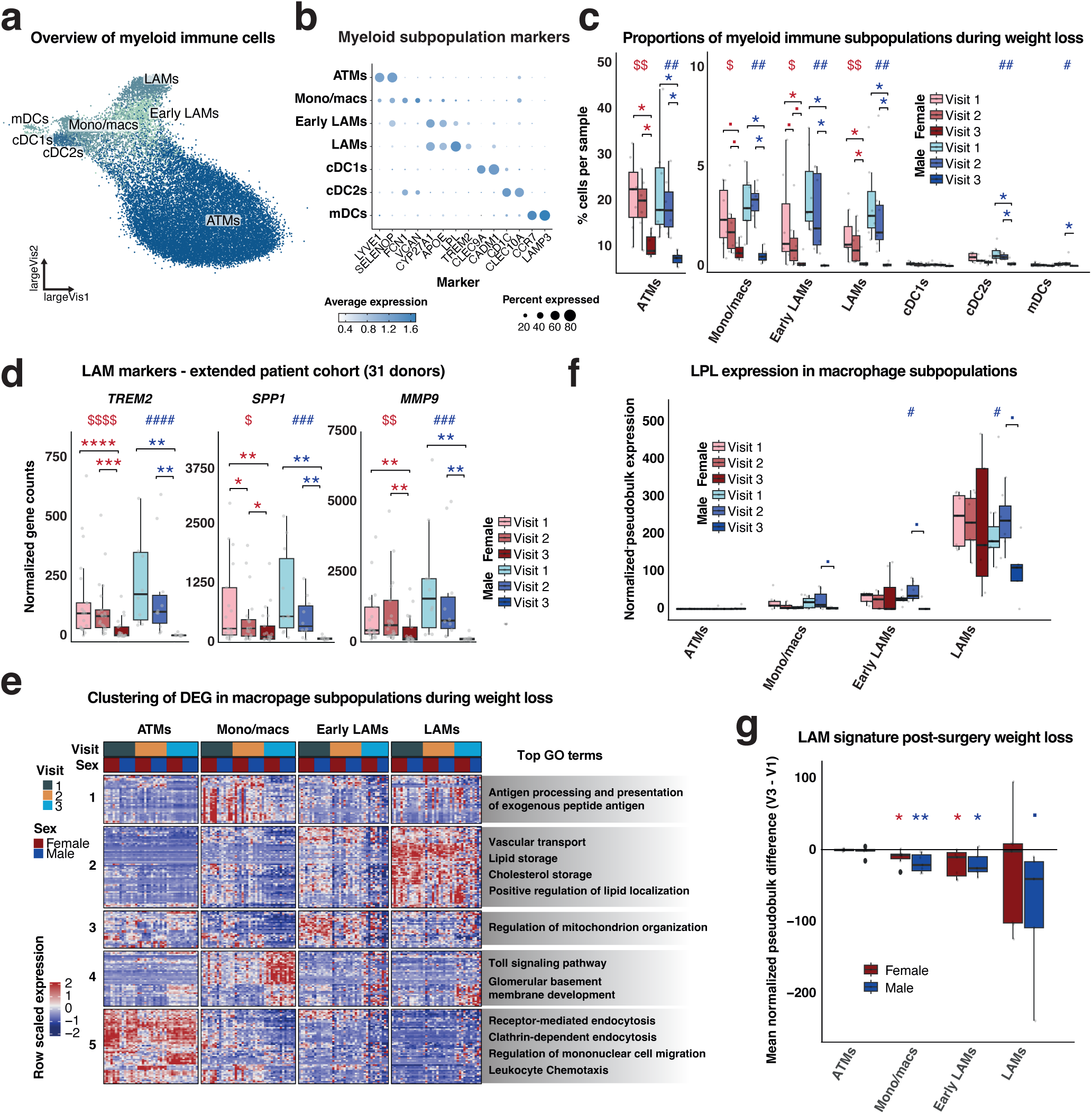
a, LargeVis representation of myeloid cells from all donors at all 3 visits. b, Marker genes for each myeloid cell population. c, The proportion of a specific myeloid immune subpopulation relative to all cells in the human ASAT dataset. d, Normalized gene counts of indicated marker genes from the bulk RNA-seq data of 31 patients (female, n=21; male, n=10) at the indicated visits (due to y-axis constraints, a few data points for *SPP1* and *MMP9* were omitted from the graph). e, Heatmap showing row-scaled expression of differentially expressed genes (DEGs) between visits in one or more of the indicated subpopulations (left) and top gene ontology (GO) term for each hierarchical gene cluster (right). f, Normalized pseudobulk expression of *LPL* from the snRNA-seq data. g, Mean normalized difference in pseudobulk signal (visit 3 – visit 1) of top LAM markers for indicated subpopulations. Box plot data: center line, median; box limits, upper and lower quartiles; whiskers, 1.5x IQR. A KW test, followed by post-hoc Wilcoxon signed-rank tests (in c and d) or Wilcoxon rank-sum test (in f) with Holm’s correction for multiple testing were performed to assess significance between visits for each sex separately. A one-sided one-sample t-test was used to determine whether the difference in normalized pseudobulk signal between visit 3 and visit 1 significantly deviated from 0 in g (▪, p<0.1; #/$/*, p<0.05; ##/$$/**, p<0.01; ###/***, p<0.001; $$$$/####/****, p<0.001). ATM, adipose tissue-resident macrophages; cDCs, conventional dendritic cells; eLAMs, (early) lipid-associated macrophages ((e)LAMs); mDCs, migratory DCs.

To investigate the transcriptional changes in macrophage subpopulations in response to weight loss, we clustered all genes significantly regulated by weight loss in one or more of the subpopulations (Fig. 3e, Extended Data Table 3). Here, cluster 1 contains genes primarily repressed by weight loss in females and linked to antigen presentation and MHC protein complex assembly (Fig. 3e and Extended Data Fig. 3l). Furthermore, the surgery-repressed cluster 2 contains several LAM marker genes linked to lipid storage (Fig. 3e-f), suggesting that both the number of LAM cells and the LAM selective gene program in macrophage subpopulations were decreased after bariatric surgery. Indeed, we found that surgery-induced weight loss led to a decrease in the expression of the top 20 LAM marker genes in several subpopulations of macrophages (Fig. 3g).

In summary, we demonstrate that all macrophage populations were dramatically reduced in ASAT after bariatric surgery and that the macrophage populations that remained in the tissue were transcriptionally reprogrammed towards a non-LAM signature.

### Lifestyle intervention in obese patients remodels the ASPC landscape

ASPCs serve diverse roles in AT, one of which is their ability to undergo adipogenesis, which is a prerequisite for maintaining healthy AT functions ^38^. ASPCs are further thought to play a key role during the development of obesity and associated complications ^39–41^. In our study, the large cluster of human ASPCs express the common marker genes, *PDGFRA* and *DCN* (Fig. 1g-h), and subclustering of these identified four distinct subpopulations (Fig. 4a-b and Extended Data Fig. 4a-c). The *DPP4*^HI^ ASPC subpopulation is marked by expression of *DPP4*, *SEMA3C* and *FBN1* and transcripts related to extracellular matrix assembly (Fig. 4b, Extended Data Table 1-2). This subcluster likely represents the human orthologous cell type of multipotent interstitial progenitor cells found in mouse AT (Extended Data Fig. 4d) ^42,43^. The *CXCL14*^HI^ ASPC subcluster is marked by transcripts related to the complement cascades and chemokine signaling, indicating that they might play a role in modulating the humoral immune response in the tissue (Extended Data Table 1-2). Based on comparisons with previously annotated mouse and human ASPCs positioned along an adipogenic differentiation trajectory ^4,8^, this cluster likely constitutes a more committed version of ASPCs than the *DPP4*^HI^ ASPCs (Extended Data Fig. 4d-e). While *PPARG*^HI^ ASPCs likely represent *bona fide* preadipocytes, as indicated by their relatively high expression levels of adipocyte marker genes like *PPARG*, *ACACB*, and *CD36* (Fig. 4b and Extended Data Fig. 4d-e) ^4^, *EPHA3^HI^*ASPCs are characterized by expression of *MEOX2, F3* and *IGFBP7*, resembling the ‘adipogenesis-regulatory’ cell (Aregs) population previously reported in mice ^40,44^.

**Fig. 4:**
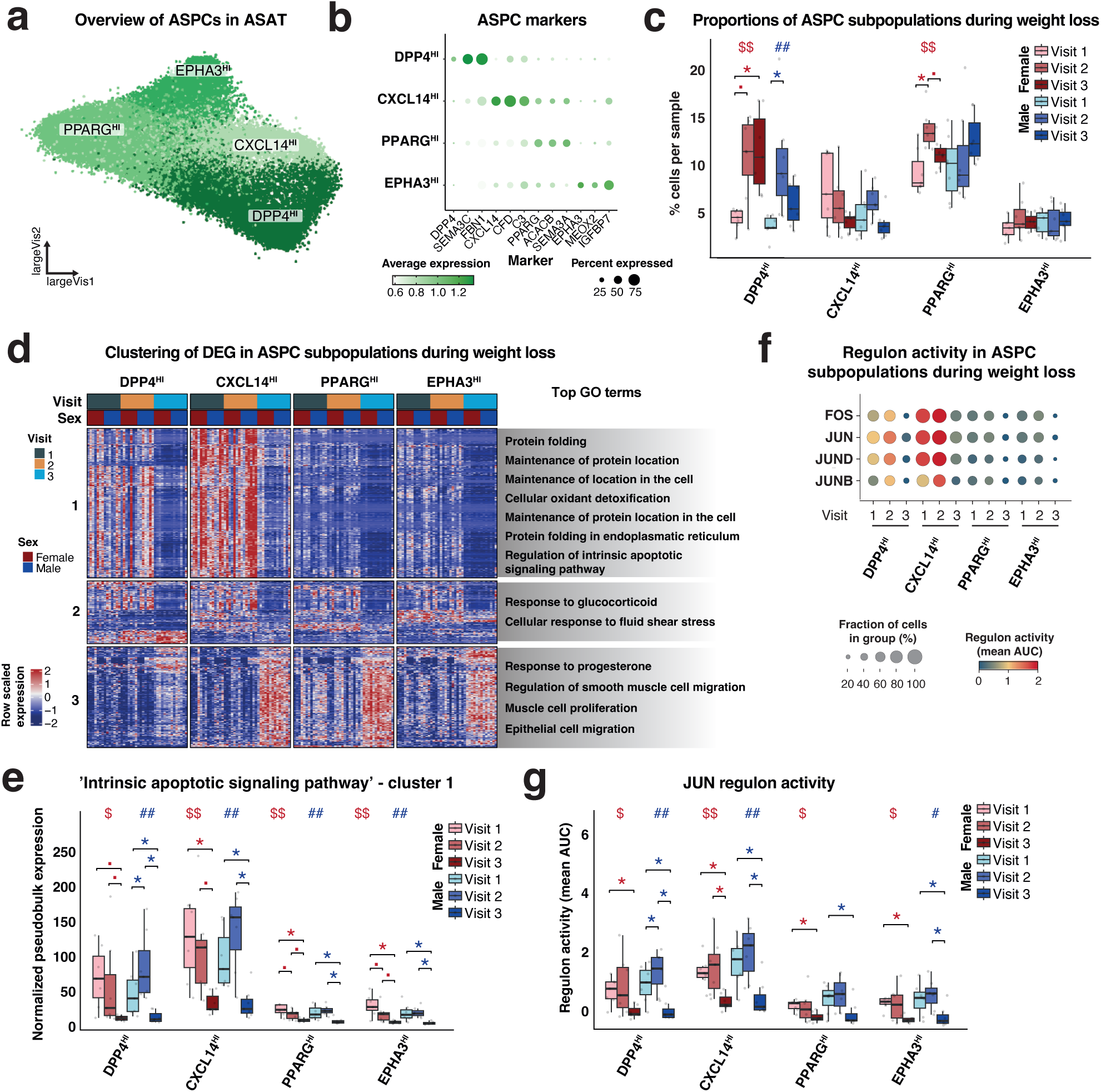
a, LargeVis representation of ASPCs from all donors at all 3 visits. b, Marker genes for each ASPC population. c, The proportion of a specific ASPC subpopulation relative to all cells in the human ASAT dataset. d, Heatmap showing row-scaled expression of DEGs between visits in one or more of the indicated subpopulations (left) and top GO term for each hierarchical gene cluster (right). e, Mean normalized pseudobulk expression of cluster 1 genes associated with the GO term GO:0097193. f, Regulon activity (mean AUC) and fraction of cells in group expressing the regulon for indicated transcription factors in each ASPC subpopulation. g, Regulon activity (mean AUC) of JUN. Box plot data: center line, median; box limits, upper and lower quartiles; whiskers, 1.5x IQR. A KW test, followed by a post-hoc Wilcoxon signed-rank test with Holm’s correction for multiple testing, was performed to assess significance between visits for each sex separately in c, e, and g (▪, p<0.1; #/$/*, p<0.05; ##/$$, p<0.01).

We and others have previously reported that obesity induces a shift in the ASPC compartment in mouse epididymal white AT (eWAT) towards more preadipocytes and fewer *DDP4+* early progenitors ^4,43^, consistent with the notion that obesity leads to increased adipogenesis in eWAT ^45^. It is currently unclear whether obesity in humans induces a similar shift towards preadipocytes. Interestingly, in the present study the proportions of early progenitors (*DPP4*^HI^ ASPCs) were increased in both sexes in response to the short-term moderate weight loss (Fig. 4c), suggesting a recruitment of these progenitors by a moderate weight loss. Moreover, the proportion of *PPARG*^HI^ ASPCs was also elevated in response to the short-term moderate weight loss in females, further indicating that the ASPC compartment in contrast to other cellular compartments in ASAT underwent an ‘early’ remodeling in response to moderate weight loss.

To investigate the transcriptional changes in the ASPC subpopulations in response to weight loss, we clustered all genes significantly regulated by weight loss in one or more of the subpopulations (Fig. 4d, Extended Data Table 4). The largest cluster (cluster 1) is primarily expressed in *DPP4*^HI^ and *CXCL14*^HI^ ASPCs and is repressed in response to surgery-induced weight loss. This cluster contains genes related to apoptotic processes and several heat shock protein family members, suggesting that these processes were less prominent in the ASPC compartment following surgery-induced weight loss (Fig. 4d-e, and Extended Data Fig. 4f). Another surgery-repressed cluster (cluster 2) contains genes related to stress and inflammatory responses, e.g., genes encoding the AP-1 TFs, *JUN*, *FOS,* and *FOSB* (Fig. 4d, Extended Data Fig. 4g, Extended Data Table 4). This prompted us to apply SCENIC ^46,47^ to investigate how the predicted regulatory networks of these TFs were affected by weight loss ^46,47^. The results indicate that the activity of the JUN, FOS, and FOSB regulons dropped markedly in all ASPC subpopulations (Fig. 4f-g and Extended Data Fig. 4h), suggesting that the significant weight loss reduced the stress and inflammatory-related activity in these ASPCs.

In summary, our data indicate that the ASPC compartment undergoes considerable compositional and transcriptional changes even after the modest lifestyle-induced weight loss, suggesting that ASPC remodeling is an early step in the healthy AT adaptation following weight loss.

### Transcriptional reprogramming of adipocyte states in response to weight loss

The adipocyte cluster constitute the third largest population of nuclei in our dataset and is marked by high expression of classical adipocyte marker genes, such as *ADIPOQ*, and *PLIN1* (Fig. 1g-h, Extended Data Fig. 5a-c). A recent integrated analyses of sxRNA-seq studies reported inconsistent adipocyte heterogeneity across different studies ^7^. Therefore, we examined the cluster stability and found that both low- and high-resolution clustering exhibited low stability of the suggested adipocyte subclusters (Extended Data Fig. 5d-f). In line with this, we found that previously proposed adipocyte subpopulation marker genes from subcutaneous and visceral human AT ^48^ did not separate well in the annotated clusters (Extended Data Fig. 5g). To identify the most polarized adipocytes in ASAT, we performed archetype analyses and selected the top 2000 cells per archetype over 10 iterations (Fig. 5a). Using the same set of predefined markers, we showed that these archetypes, *DGAT2*^Arc^, *CLSTN2*^Arc^, and *PRSS23*^Arc^, are more transcriptionally distinct than the annotated subpopulations, indicating the presence of extreme ‘archetypical’ adipocyte transcriptional states in ASAT (Fig. 5b). *DGAT2*^Arc^ cells express high levels of several lipogenic markers such as *LPL, DGAT2, CD36* and *ADIPOQ* and are enriched for transcripts related to fatty acid metabolic processes (Extended Data Table 1-2). This cell state therefore likely resembles previously described ‘lipogenic’ adipocytes, assumed to be adipocytes with a high capacity for *de novo* lipogenesis ^4,5^. *CLSTN2*^Arc^ cells express some markers related to inflammatory signaling, such as *CBLB*, whereas *PRSS23*^Arc^ cells express high levels of genes linked to cellular signaling, e.g., *CAV2*. Differential expression analyses identified 4 clusters of genes with distinct expression patterns in response to weight loss in the 3 archetypes (Fig. 5c). Three of these clusters contain genes induced in response to surgery-induced weight loss to various extents in the adipocyte archetypes, whereas the last cluster contains genes downregulated by surgery. One cluster (cluster 2) consists of genes upregulated across all adipocyte archetypes, which are linked to beta oxidation as well as lipid and glucose metabolic processes, such as *DECR*, *LIPA*, and *PLIN5* (Fig. 5d, Extended Data Table 5). Another cluster (cluster 3) contains genes primarily induced in *CSLTN2*^Arc^ adipocytes in response to weight loss, and these genes are associated with signaling processes, including *MAP3K5, and AKT3* (Extended Data Fig. 5h). Finally, the downregulated cluster, cluster 4, contains several collagen- and matrix metalloproteinases-associated genes (Extended Data Fig. 5i and Extended Data Table 5), indicating that extracellular matrix remodeling associated with adipocytes is affected by weight loss.

**Fig. 5:**
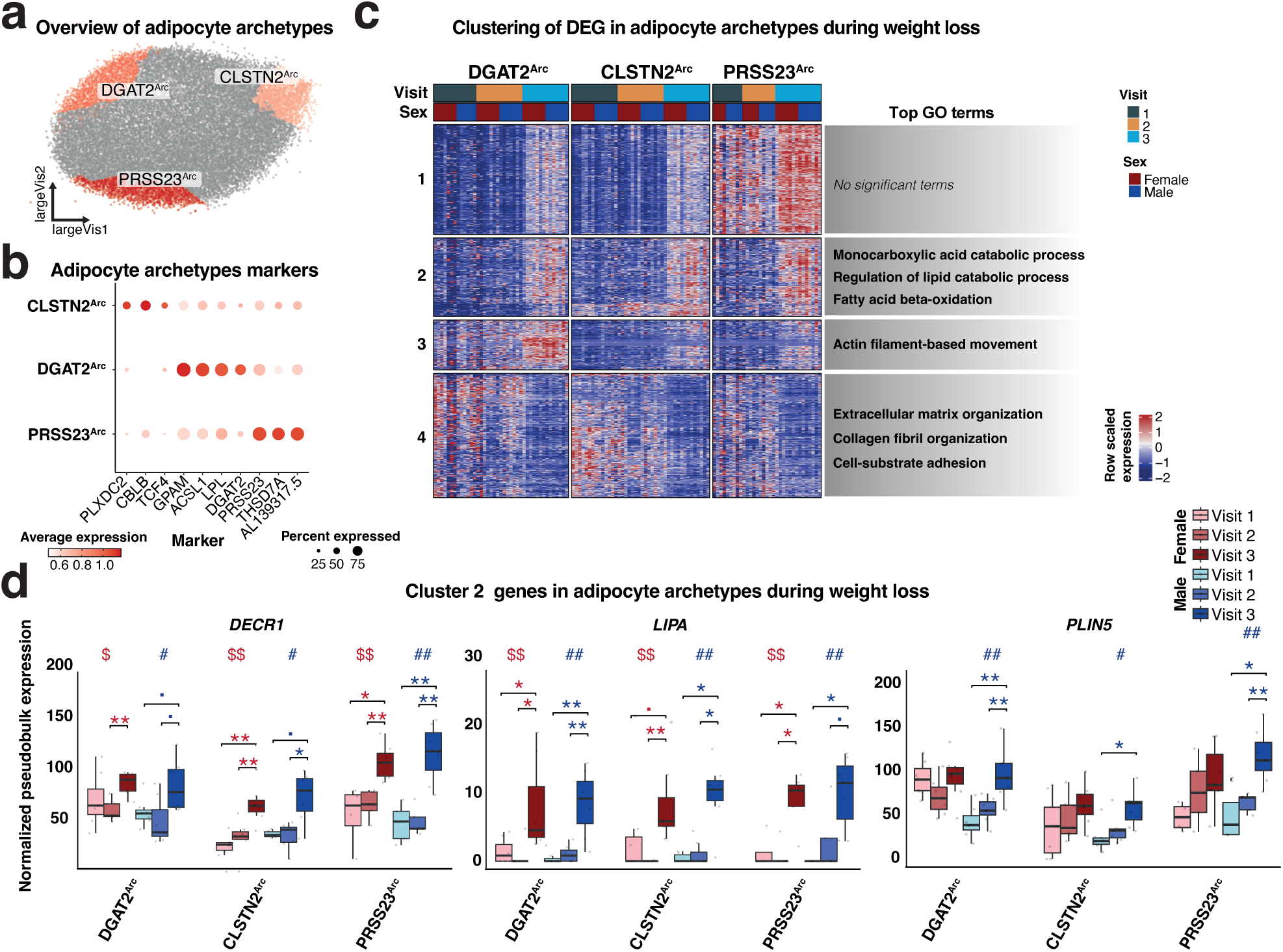
a, LargeVis representation of adipocytes and adipocyte archetypes from all donors at all 3 visits. b, Marker genes for each adipocyte archetypes. c, Heatmap showing row-scaled expression of DEGs between visits in one or more of the indicated archetypes (left) and top GO term for each hierarchical gene cluster (right). d, Mean normalized pseudobulk expression of indicated genes from cluster 2. Box plot data: center line, median; box limits, upper and lower quartiles; whiskers, 1.5x IQR. A KW test, followed by a post-hoc Wilcoxon rank-sum test with Holm’s correction for multiple testing, was performed to assess significance between visits for each sex separately in d (▪, p<0.1; #/$/*, p<0.05; ##/$$/**, p<0.01).

In summary, this study did not provide evidence of highly distinct adipocyte subpopulations in human ASAT. Nonetheless, we found evidence for the existence of transcriptional extreme states of adipocytes that undergo a slightly distinct transcriptional reprogramming following surgery-induced weight loss. Overall, bariatric surgery induced an increased lipogenic oxidative signature and a decreased extracellular matrix remodeling profile in adipocytes.

### A map of human *in vivo* adipogenesis

Since we captured both ASPCs and mature adipocytes in our snRNA-seq approach, we asked whether we could establish an ASPC differentiation hierarchy and trace human *in vivo* adipogenesis. Due to the transcriptional similarity between potential adipocyte subtypes, we chose to analyze the mature adipocytes as one population. Our initial analyses of the transcriptional profile indicated that *DPP4*^HI^ ASPCs represented multipotent progenitors, whereas *PPARG*^HI^ ASPCs were committed preadipocytes. To explore this further, we mapped the trajectories between ASPCs and mature adipocytes using diffusion maps and pseudo-time analyses ^49,50^. This identified two distinct trajectories with one trajectory likely representing adipogenesis and one trajectory representing an alternative specification for the ASPCs (Fig. 6a). The adipogenesis branch appears to be initiated in *DPP4*^HI^ ASPCs and progress through first *CXCL14^HI^*, then *PPARG*^HI^ ASPCs before ultimately leading to the formation of mature adipocytes (Fig. 6a-b). Interestingly, *EPHA3^HI^* ASPCs does not appear to be part of the adipogenesis trajectory (Fig. 6a), thereby supporting that these may have an alternative, regulatory role in the tissue, as previously suggested in mice ^40,44^.

**Fig. 6:**
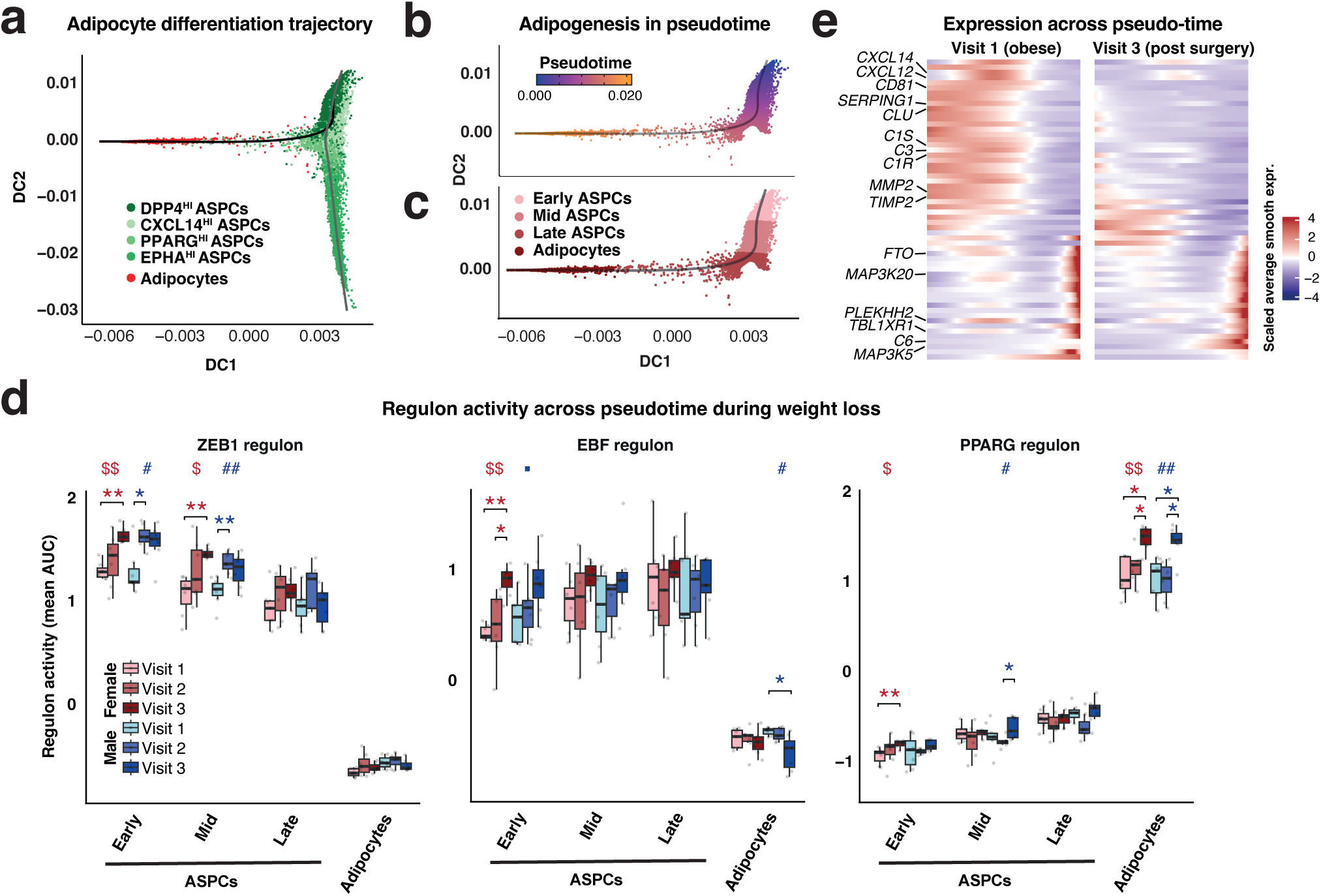
a, Diffusion maps of ASPC and adipocyte nuclei with indication of two distinct trajectories. b-c, Diffusion maps of ASPC and adipocyte nuclei belonging to the adipogenic differentiation trajectory with indication of pseudotime (b) or with re-classification of nuclei in accordance with their differentiation state (c). d, Regulon activity (mean AUC) of indicated transcription factors at different cellular stages during adipocyte differentiation (as defined in b-c). e, Heatmap showing average scaled expressions of genes differentially expressed between visit 1 and 3 and along pseudo-time across the trajectory (in b) for visit 1 and visit 3. Box plot data: center line, median; box limits, upper and lower quartiles; whiskers, 1.5x IQR. A KW test, followed by post-hoc Wilcoxon rank-sum test with Holm’s correction for multiple testing, was performed to assess significance between visits for each sex separately in d (▪, p<0.1; #/$/*, p<0.05; ##/$$/**, p<0.01).

To further explore the human adipogenesis trajectory, we identified 240 genes that were significantly differentially expressed across this trajectory (Extended Data Fig. 6a). We clustered these genes based on their temporal expression patterns and confirmed that genes expressed early in this trajectory were related to extracellular matrix organization, whereas genes expressed late in the trajectory can be associated to lipid metabolic processes (Extended Data Fig. 6b, Extended Data Table 6). 14 of these genes encode transcriptional regulators, some of which are known for their role in adipogenesis (Extended Data Fig. 6c). This includes the master adipogenic regulator, PPARγ and the early factors, early B-cell factor 2 (EBF2) and zinc finger E-box-binding homeobox 1 (ZEB1), which have been shown to induce adipogenic competency in mouse preadipocytes ^51,52^. To examine how weight loss affects these adipogenic TFs and their regulatory networks across the adipogenic trajectory, we divided the adipogenic trajectory into four bins, representing early, mid, and late preadipocytes, as well as mature adipocytes (Fig. 6c). While *ZEB1* expression was not affected by weight loss in any of the preadipocyte states (Extended Data Fig. 6d), the activity of the ZEB1 regulon was markedly increased already following lifestyle-induced weight loss in the early and mid-preadipocytes in males (and trending in females) (Fig. 6d). This suggests that ZEB1 may be post-transcriptionally activated and playing a role in the early adaptation of adipogenesis following weight loss. The activity of the EBF regulon was also induced by weight loss in early preadipocytes, although only after more substantial weight loss (Fig. 6d), suggesting that EBF2 could play a role in promoting healthy adipogenesis in the post-obese, lean state. Additionally, the PPARG regulon showed markedly increased activity in adipocytes following bariatric surgery in both sexes (Fig. 6d), suggesting improved transcriptional regulation of the adipogenic program after significant weight loss. In females, the PPARG regulon was also induced in early ASPCs following surgery-induced weight loss, further supporting an increased adipogenic potential.

Finally, we focused on the specific genes differentially regulated in the pseudotime-space in the obese and the post-surgery state. In the obese state, several genes linked to complement activation and pro-inflammatory processes, such as *C3*, *C1R*, *C1S*, *CXCL12*, and *CXCL14,* were expressed at high levels during adipogenesis (Fig. 6e), which is not observed in the post-obese lean state. This suggests an abnormal temporal regulation of these genes during adipogenesis in obesity, possibly affecting the ability of the preadipocytes to undergo adipogenesis, thereby limiting healthy expansion of the ASAT.

### Altered intercellular communication between ASPCs and adipocytes in response to weight loss

Recent studies have suggested that the cellular crosstalk between adipocyte cell types is significantly altered in obesity ^4,5^; however, so far there is little insight into how this crosstalk may be affected by weight loss. To assess potential interactions between the identified cell types in ASAT, we computed a cellular communication and interaction (CCI) score using LIANA as a proxy for ligand-receptor interactions in a given condition ^53^. In the obese state, ASPCs, adipocytes and endothelial cells were predicted as the main interacting cell types (Fig. 7a). Upon surgery-induced weight loss, interaction scores between most cell types were decreased, suggesting less intercellular crosstalk after weight loss (Fig. 7a). To further explore the changes in cellular crosstalk in response to different weight loss modalities, we built a communication tensor containing the 7 major cell types in AT ^54^. By decomposing the tensor from our datasets into 11 factors, we associated communication patterns between the different AT cell types to different modes of weight loss (Fig. 7b + Extended Data Fig. 7a). Most notably, factor 3 represents a communication pattern in which the cellular interactions are reduced following surgery-induced weight loss (Fig. 7b). Factor 3 captures cellular interactions originating from ASPCs to various other AT cell types, including adipocytes, vascular cells, and immune cells (Fig. 7c + Extended Data Fig. 7b-c). The key ligand-receptor interactions in factor 3 (Fig. 7d) involve several ASPC ligands, including the complement factor C3, which previously have been linked to inflammation and insulin resistance in AT ^55,56^. The interactions of these ligands with receptors on other adipose cell types are predicted to decrease following weight loss (Fig. 7d). This may be attributed to the lack of transcriptional induction of genes encoding these ligands during adipogenesis after weight loss (Fig. 6e and 7e).

**Fig. 7:**
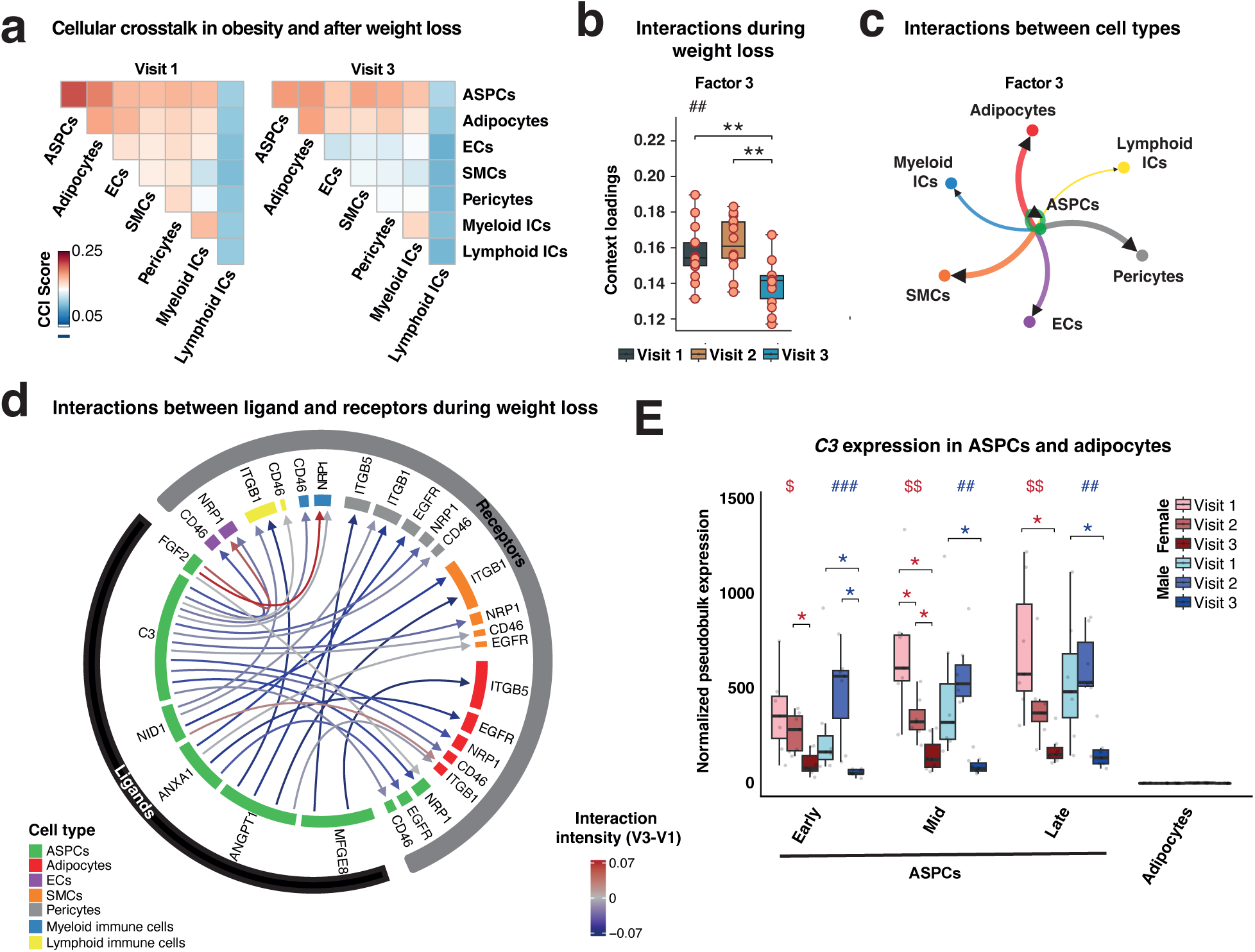
a, Heat map showing the cellular communication and interaction (CCI) score between different cell types for visit 1 and visit 3 as determined by LIANA. b, Sample loadings for each visit for factor 3 as determined by LIANA and Tensor-cell2cell. c, Interaction map illustrating cell types associated with factor 3. The arrowhead indicates the direction of the cross talk (sender → receiver) and the line width indicates the strength of interaction between each cell type. d, Circle plot showing the key factor 3-enriched interaction links between ASPC ligands and their target receptor on the indicated cell types. The arrowhead indicates the direction of the cross talk (sender → receiver) and the line color indicates the difference in interaction intensity between visit 3 and visit 1. e, Mean normalized pseudobulk expression of *C3* from the snRNA-seq data. Box plot data: center line, median; box limits, upper and lower quartiles; whiskers, 1.5x IQR. A KW test, followed by a post-hoc Wilcoxon signed-ranked test with Holm’s correction for multiple testing, was performed to assess significance between visits for each sex separately in b, and e ($/*, p<0.05; ##/$$/**, p<0.01; ###, p<0.001).

Taken together, our findings propose a model where the reduction of multiple pro-inflammatory ligands in ASPCs in the post-obese, lean state likely diminishes the crosstalk between ASPCs and neighboring cells. This dampened signaling may create a healthier microenvironment, which in turn promotes favorable AT remodeling.

## Discussion

Here we present a single cell-resolved map of the cellular and molecular plasticity of human ASAT in men and women in response to both a moderate lifestyle-induced weight loss as well as to a more dramatic bariatric surgery-induced weight loss. While lifestyle-induced weight loss led to moderate compositional and transcriptional changes in a few cellular subpopulations, bariatric surgery-induced weight loss resulted in profound changes in both composition and gene expression in both females and males. Overall, these changes indicate that two-years post-surgery there was not only significantly less inflammation in ASAT but also increased vascularization and increased adipogenic potential.

Several studies in mouse AT have shown that angiogenesis is tightly associated with AT growth ^57–59^. Moreover, single cell studies in humans have demonstrated that the proportions of vascular cells in AT correlate negatively with BMI and that obesity leads to an impairment of the angiogenic response in specific vascular subpopulations ^5,31^. Our data suggest that bariatric surgery can reverse at least some of these obesity-related changes in the vascular network, consistent with recent findings in mice ^32^. We show that bariatric surgery resulted in a striking overall increase in the proportions of ECs, with a specific expansion of vECs and cECs. While the increase in vECs may primarily be a result of structural changes in the overall vascular network, it is likely that the elevated proportion of cells within the capillary beds results from greater proliferative and migratory abilities of cECs ^60^. Our data therefore supports that sustained weight loss via cEC remodeling promotes angiogenesis to ensure adequate oxygenation and nutrient supply to the remaining adipocytes and stromal cells, facilitating healthy AT remodeling.

Flow cytometry and single cell sequencing studies have shown that myeloid cells can comprise up to 50% of all cells in obese mice ^4,36,37^. A similar, albeit less dramatic, increase in myeloid cells has been reported in human obesity ^35^. Studies on weight loss show substantial variability in findings related to reversal of AT inflammation, potentially due to differences in weight loss regimens, specific fat depots examined, durations of interventions, and levels of obesity severity. Several studies in mice have shown that short-term weight loss results in an initial increase in macrophage infiltration in AT ^10,11^. In contrast, studies in humans report mixed outcomes, with lifestyle-induced short-term weight loss associated with increased, decreased, or unchanged AT inflammation ^12–16,25^. The reports on the effects of dramatic and sustained weight loss results are also conflicting. Several studies in mice have indicated that while inflammation is reduced, inflammatory signals persist in AT, even after dramatic and sustained weight loss ^10,11,61^. In humans, a profound inflammatory memory has been reported to persist as long as 2 years after bariatric surgery as determined by qPCR of inflammatory transcripts and detection of CLS ^18^, microarray analysis of bulk RNA ^62^, and most recently by snRNA-seq ^25^. By contrast, other studies report a prominent reduction of inflammatory markers in ASAT following bariatric surgery based on qPCR ^63,64^ and immunohistology ^22^.

In the present study, we applied snRNA-seq, bulk RNA-seq and 3D light microscopy to carefully investigate ASAT remodeling during both modest lifestyle-induced and dramatic surgery-induced weight loss. We found that following a ∼10% lifestyle-induced weight loss, the proportions of macrophage subpopulations remained relatively stable, suggesting that macrophage lipid handling continued to play a role in supporting healthy AT remodeling during the early stages of weight loss.^10,11,13,14,65^. Interestingly, however, there was a clear trend towards a decrease in CLS, indicating a modest improvement in AT health following moderate weight loss. Following the surgery-induced weight loss, the proportions of myeloid immune cells and CLS numbers were dramatically decreased. Although this reduction was evident across all macrophage subpopulations, the clearance of LAMs from ASAT following weight loss was particularly remarkable. Furthermore, macrophages that remained in ASAT seemed to suppress both their inflammatory and lipid handling gene programs two years post-surgery. Thus, our data suggest that, in a post-obese lean state, macrophage-mediated lipid handling does not play a prominent role in maintaining ASAT homeostasis.

AT hyperplasia, characterized morphologically by the presence of many small adipocytes, has been linked to protection against obesity-related complications ^66,67^. The hyperplastic potential of AT likely depends on both the quantity and quality (i.e., proliferative capacity and ability to undergo adipogenesis) of ASPCs, which is believed to be compromised in obesity ^23,68^. In our study, we showed that the relative proportions of DPP4^HI^ ASPCs in ASAT in both sexes as well as the relative proportion of PPARG^HI^ ASPCs in females were elevated already in the early phases of a weight loss. Although we have not formally demonstrated that the increase in specific ASPC subpopulations would restore a normal hyperplastic potential in the tissue, there are several indications supporting this notion. First in mice, DPP4+ progenitors in both SAT and VAT have been shown to contribute to adipogenesis ^42,69^, which would suggest that the increase in pro-adipogenic ASPCs induced by weight loss in humans may enhance adipogenic potential in ASAT. Secondly, our data supports that regulatory networks controlled by transcriptional regulators that have previously been shown to be proadipogenic (e.g., EBF2, and ZEB1) ^51,52^ are more active in the early phases of adipogenesis after weight loss. Third, the elevation of PPARG regulon activity in the late stage adipogenesis after weight loss would support the transitioning of preadipocytes into fully mature, lipid-laden adipocytes that are functionally prepared to store fat and secrete adipokines. The elevated activity of PPARG would further ensure a more efficient lipid metabolism and enhanced insulin responsiveness in mature adipocytes, thereby stimulating adipocyte metabolic function. Interestingly, a recent snRNA-seq study of human WAT reported that many key metabolic genes in adipocytes remain downregulated even 2 years after bariatric surgery, suggesting that weight loss are not sufficient to resolve obesogenic-induced changes in adipocytes ^25^. In contrast, in the present study we found that genes associated with lipid and glucose metabolic processes were markedly induced in adipocytes 2 years post bariatric surgery, which would align with the elevated PPARG activity. As our study did not include lean individuals, we cannot determine whether the changes in metabolic genes are fully resolved or only partially improved. Nonetheless, our findings demonstrate a marked a shift towards a much healthier preadipocyte and adipocyte state following surgery-induced weight loss.

In addition to the elevated activity of pro-adipogenic factors during adipogenesis, our analyses also reveal that surgery-induced weight loss suppresses obesity-induced genes in preadipocytes, some of which have been linked to development of insulin resistance and increased macrophage infiltration of AT ^55,56^. Some of these genes encode complement factors that are predicted to be crucial signaling molecules secreted from ASPCs, which contribute to the exacerbation of inflammation in AT by directly interacting with immune cells and adipocytes in the obese state. Following surgery-induced weight loss, complement factors, such as C3, are less strongly induced during adipogenesis, resulting in less pro-inflammatory crosstalk between ASPCs and other cell types in AT. This further supports the notion of healthy remodeling of AT following weight loss.

In conclusion, we report that bariatric surgery-induced weight loss leads to major compositional, structural and transcriptional remodeling in ASAT, with increasing vascularization and reducing inflammation, while modest weight loss promotes progenitors and proadipogenic genes, aiding early metabolic improvements. Future studies should investigate the systemic as well as local mechanisms involved in driving these dramatic changes in the ASAT during weight loss and determine the role of these changes in driving the improvement of cardiometabolic health.

## Methods

### Study design and participants

The research described in this manuscript complies with all relevant ethical regulations. Written informed consent was obtained from eligible participants before any study-related procedures. ASAT, liver biopsies, and blood samples were obtained from participants enrolled in an ongoing prospective interventional case-control study, PROMETHEUS. This study is a liver biopsy-controlled, single-center study conducted in Denmark. The inclusion criteria include participants aged 18-70 years with a BMI of ≥35 kg/m². Exclusion criteria include excessive alcohol consumption (>12 g/day for women and >24 g/day for men), the presence of other known or newly discovered chronic liver diseases, use of hepatotoxic medications (such as glucocorticoids, tamoxifen, or amiodarone), short life expectancy, or contraindications for liver biopsy. For the present study we included male (10 subjects) and female subjects (21 subjects) scheduled for bariatric surgery and biopsies were taken at the time of enrollment (visit 1), at the time of bariatric surgery following a mandatory lifestyle-induced weight loss (visit 2), and then approximately two-years post-surgery (visit 3). We selected 7 males and 7 females with the high MetS score at visit 1 for single cell analyses.

PROMETHEUS is registered with OPEN.rsyd.dk (OP-551, Odense Patient data Explorative Network) and ClinicalTrials.gov (NCT03535142). The study was approved by the regional committee on health research ethics, and all participants provided written informed consent (S-20170210) prior to participation. Study data, including biometric information (e.g., height, weight, and BMI), anthropometric measurements, and pharmacological treatment data, were collected prospectively and managed using REDCap (Research Electronic Data Capture) tools hosted at OPEN.rsyd.dk. REDCap is a secure web-based software platform designed to support data capture for research studies (https://www.sdu.dk/en/forskning/open).

### Sampling of human ASAT samples

ASAT biopsies were sampled under sterile conditions by two trained clinicians after a bedside ultrasound guided marking for best placement of a liver biopsy at visit 1 and 3 with the same protocol. The ASAT biopsy site was therefore typically between the seventh and eighth intercostal space at the mid-axillary line. Samples from visit 2 were taken during surgery, and through the central trocar port-site, mid-abdominal wall and few centimeters above the umbilicus. Samples were immediately released into sterile saline water, divided into smaller pieces and preserved immediately in RNAlater (Sigma-Aldrich, St. Louis, MO) or snap frozen in liquid nitrogen. Blood samples were collected by an experienced laboratory technician. Biochemical analyses were conducted following standard regional protocols using commercially available kits. All samples were processed by specialized research biochemical technicians and stored at −80 °C.

### Histology

#### Adipose Tissue Clearing and Immunofluorescent Staining

WAT histology was performed using the Adipo-Clear protocol ^70^. Cleared tissues were stained with primary antibodies (Rabbit anti-PLIN1 (Abcam, ab3526) and Rat anti-F4/80 (Invitrogen, 14-4801-85)) and secondary antibodies (Donkey anti-Rabbit Alexa FluorTM 488 (Invitrogen, A-21206) and Donkey anti-Rat Alexa FluorTM 568 (Invitrogen, A78946)) diluted 1:500 for 5 days each, and 0.1 µg/mL DAPI (Sigma Aldrich, D9542) for the final day of secondary incubation. 3D imaging was conducted using light-sheet fluorescence microscopy (UltraMicroscope BlazeTM, Miltenyi Biotec) with MACS Imaging Solution (Miltenyi Biotec, 130-128-511). Imaging parameters included dual-sided triple sheet illumination, channels for DAPI (405/30 nm excitation, 460/40 nm emission), PLIN1 (450/20 nm excitation, 595/40 nm emission), and F4/80 (560/40 nm excitation, 620/60 nm emission), 10x magnification, 1000 µm stacks (2 µm step size), autofocus, and dynamic horizontal focus. Representative images were captured using confocal fluorescence microscopy (Nikon Eclipse Ti2 Inverted Confocal A1) with channels for DAPI (407 nm excitation, 425-475 nm emission), PLIN1 (488 nm excitation, 500-550 nm emission), and F4/80 (561 nm excitation, 570-620 nm emission). Settings included a 10x objective, 1.2 airy unit pinhole, and 16x averaging.

### RNA extraction

ASAT was homogenized in TRI Reagent® (Sigma-Aldrich) using the FastPrep-24™ instrument (MP biomedicals) followed by RNA extraction using EconoSpin columns (Epoch). NEBNext Ultra RNA Library Prep Kit for Illumina (New England Biolabs, San Diego, CA) was used for construction of libraries according to the manufacturer’s protocol. RNA was paired-end-sequenced using the NovaSeq™ 6000 platform (Illumina, San Diego, CA).

### Nuclei Isolation

Nuclei isolation from ASAT biopsies from visit 1 and 2 was performed essentially as previously described ^71^. For visit 3 biopsies, the minced tissue was homogenized in 2 mL of Nuclei Extraction Buffer (Miltenyi Biotec, 130-128-024) containing 0.5 U/mL RNase inhibitor (New England Biolabs, M0314) using the 4C_nuclei_1 program on the gentleMACS^TM^ Octo Dissociator (Miltenyi Biotec, 130-096-427) with gentleMACS^TM^ C Tubes (Miltenyi Biotec, 130-093-237). The nuclei suspension was then filtered through a MACS® SmartStrainer (70 µm) and subsequently centrifuged at 500×g at 4 °C for 10 min. The nuclei pellet was resuspended in 100 µL nuclei resuspension buffer [1XPBS in DEPC H_2_O, 1 mg/mL BSA (Sigma, B6917), and 0.04 U/μL RNase inhibitor. All solutions were sterile filtered prior to use. All nuclei suspension was filtered through a pre-wetted 40 µm Flowmi® Cell Strainer (Merck, BAH136800070), and nuclei were counted using a Bürker pattern counting chamber (Sigma-Aldrich, BR718920) and Typan Blue staining (Bio-Rad, 1450022). An equal number of nuclei from 7 samples were randomly pooled into 6 pools.

### Single-Nucleus RNA-Sequencing

Immediately following nuclei isolation, 20,000 nuclei were loaded onto the 10x Genomics Chromium controller (10X Genomics, PN110203). Libraries were prepared following the manufacturer’s instructions using the 10x Genomics Chromium Next GEM Single Cell 3’ Reagents Kits v3.1 (Dual Index) (User Guide CG000315 Rev E) [Next GEM Single Cell 3’ Gel Beads v3.1 (10X Genomics, PN-2000164), Next GEM Chip G (10X Genomics, PN-2000177), Chromium Next GEM Single Cell 3’ GEM Kit v3.1 (10X Genomics, PN-1000123), Library Construction Kit (10X Genomics, PN-1000190), Dual Index Kit TT Set A (10X Genomics, PN-1000215)]. Sequencing was performed on an Illumina NovaSeq 6000 System (10X Genomics, 20012850), targeting approximately 50,000 raw reads per nucleus.

### Quantification and Statistical Analysis

Clinical data was assessed for normality using the Shapiro-Wilk and Kolmogorov-Smirnov tests. Parametric tests were applied to compare groups if the data satisfied the normality assumption (p > 0.05); otherwise, non-parametric tests were used as indicated. Outliers were identified using the ROUT method with a Q value of 1%. Sequencing data, due to its typical distribution patterns, was assumed to be non-normally distributed, and non-parametric tests were consistently employed for group comparisons.

#### 3D image analysis

Light-sheet images were analyzed using Arivis Pro (v4.2.1). Adipocyte size (>1000 adipocytes per sample) was quantified from PLIN1 signals using a pipeline with denoising (median filter, 2.5 µm), membrane-based segmentation (170-270 min intensity, 20-50% split sensitivity, 200 µm max diameter), and object filtration (touching edge filter, volume filter 8,181-4,188,790 µm³ corresponding to 25-200 µm diameter, sphericity filter 0.4-1.0). CLS density was determined by counting F4/80-positive CLSs, normalized to tissue volume quantified via PLIN1 signal analysis with intensity threshold segmentation and volume filtering (>1,000,000 µm³). Representative confocal images were processed with Denoise.AI (NIS-Elements AR, v5.30.02).

#### Bulk RNA-seq data analysis

Quality of raw sequencing was assessed using *FastQC* (v.0.11.9) ^72^ and *MultiQC* (v1.10.1) ^73^. Raw bulk RNA-seq data were aligned to the human genome assembly (GRCh38, Ensembl release 101) using *STAR* (version 2.7.11a) ^74^. Aligned reads were sorted and indexed using *Samtools* (version 1.17) ^75^ and low low-quality reads were removed using MAPQ < 30, SAM flag = 780. Exon reads were quantified using *FeatureCounts* (version 2.0.6) ^76^.

#### Single nucleus RNA-seq

Raw snRNA-seq data were aligned using 10x Genomics *Cell Ranger* v. 7.1.0 ^77^ with their GRCh38-2020-A reference. Samples were demultiplexed based on bulk RNA sequencing samples from the donors included here. In short, bulk or single-nucleus sequencing data were aligned using *STAR* ^74^. Valid single-nucleus barcodes were identified using *valiDrops* ^78^. Donor SNPs were identified using C*ellsnp-lite* ^79^. Demultiplexing was performed using *vireo* ^80^. Filtering and QC was performed using *CRMetrics* ^81^ with depth = 1,000. We removed mitochondrial and ribosomal gene counts from our count matrices. Within *CRMetrics*, we adjusted for ambient RNA using *CellBender* ^82^. Doublets were identified using *DoubletDetection* ^83^. Next, we normalized counts using *Pagoda2* ^84^ before integrating and embedding all samples using *Conos* ^26^. Differentially expressed genes were identified using *Cacoa* ^85^. RNA velocity estimations were carried out using *velocyto* ^86^ for counting spliced/unspliced reads followed by analyses in *cellDancer* ^87^ and *Dynamo* ^88^ in combination. *LIANA+* ^53^ was used to perform cell-cell interaction analyses, and *LIANA* and *Tensor-cell2cell* ^89^ were combined to decipher cell-cell communication across multiple samples ^54^. Predicted cellular interactions were plotted using *circlize* ^90^.

Regulons were identified using *SCENIC* ^46,47^. Gene ontology analyses were performed using *clusterProfiler* and plotted using *enrichPlot* ^91^. Statistics were calculated and added to plots using a combination of *ggpubr* ^92^ and *rstatix* ^93^.

Pseudotime trajectories were estimated using *slingshot* ^49^ based on diffusion map embeddings made with a combination of *destiny* ^50^, *Seurat* ^94^ and *harmony* ^95^. To infer pseudotime-relevant genes, using *gamm4* ^96^, we fitted three generalized additive mixed models to each gene with donor as a random variable: a null model where gene expression (GEX) ∼ 1, a pseudotime model where GEX ∼ pseudotime, and an interaction model where GEX ∼ pseudotime + visit. We stratified genes significantly explained by the models dependent on what we wanted to investigate, i.e. we selected genes significantly described by pseudotime but not by the interaction, genes described by the interaction, and genes better described by the interaction over pseudotime. Then, we fitted and smoothed the GEX based on pseudotime using *tradeSeq* ^97^ before plotting.

Most plots were made with *ggplot2* ^98^, and heatmaps were plotted with *ComplexHeatmap* ^99^.

#### Comparison with publicly available data sets

Cell type and subpopulation similarity were assessed by comparing our annotations to human annotations in WATLAS ^8^. Gene expression counts were log-transformed and averaged per cell type to create pseudobulk profiles. Pairwise Pearson correlations were then calculated for all cell types using the subset of shared genes between the two datasets. To compare human ASPCs with their mouse counterparts, marker genes from significant fibro-adipogenic subpopulations identified in ^4^ were used as gene sets. Mouse gene symbols were mapped to their human orthologs using the *mygene* ^100^. Gene set scores were then calculated using *scanpy* ^101^ with a minimum control size of 50.

## Data and code availability

Processed snRNA-seq data will be made available upon publication via the Single Cell Portal (SCP2849). Reproducible code for figures in this manuscript is available at Github (https://github.com/rrydbirk/weight-loss-study).

## Acknowledgments

This work was supported by grants from the Danish National Research Foundation to the Center for Functional Genomics and Tissue Plasticity (ATLAS) (Project grant: 141); the Novo Nordisk Foundation (NNF18OC0033444 Challenge Grant to the Center for Adipocyte Signaling; NNF21SA0072102); the Lundbeck Foundation (R413-2022-471).

## Author information

### Authors and Affiliations

**Center for Functional Genomics and Tissue Plasticity (ATLAS), Department of Biochemistry and Molecular Biology, University of Southern Denmark (SDU), Odense, Denmark**

Anne Loft, Rasmus Rydbirk, Ellen Gammelmark Klinggaard, Elvira Laila Van Hauwaert, Andreas Fønns Møller, Trine Vestergaard Dam, Babukrishna Maniyadath, Ronni Nielsen, Søren Fisker Schmidt, Jesper Grud Skat Madsen, Susanne Mandrup.

**ATLAS, Department of Gastroenterology and Hepatology, University Hospital of Southern Denmark, Esbjerg, Denmark.**

Charlotte Wilhelmina Wernberg, Mette Enok Munk Lauridsen

**ATLAS, Center for Liver Research (FLASH), Department of Gastroenterology and Hepatology, Odense University Hospital, Odense, Denmark.**

Aleksander Krag

**The Novo Nordisk Foundation Center for Genomic Mechanisms of Disease, Broad Institute of MIT and Harvard, Cambridge, MA, 02142, USA**

Andreas Fønns Møller, Jesper Grud Skat Madsen, Susanne Mandrup

**Department of Biomedicine, Aarhus University, Aarhus, 8000, Denmark; Steno Diabetes Center Aarhus, Aarhus University Hospital, Aarhus, Denmark**

Mohamed Nabil Hassan, Joanna Kalucka

### Contributions

Conceptualization and study design were carried out by A.L., R.R., E.G.K., E.L.V.H., J.G.S.M., and S.M. Acquisition of data was carried out by E.L.V.H., E.G.K., C.W.W., T.V.D., B.M., R.N., A.K., M.E.M.L. Analysis and interpretation of data were carried out by A.L., R.R., E.G.K., E.L.V.H., A.F.M., M.N.H., J.K., S.F.S., J.G.S.M., and S.M.

### Corresponding authors

Correspondence to Anne Loft, Jesper Grud Skat Madsen, and Susanne Mandrup.

