## Extended Data Figures for "Single cell-resolved transcriptional dynamics of human subcutaneous adipose tissue during lifestyle- and bariatric surgery-induced weight loss"

### Extended Data Fig.1

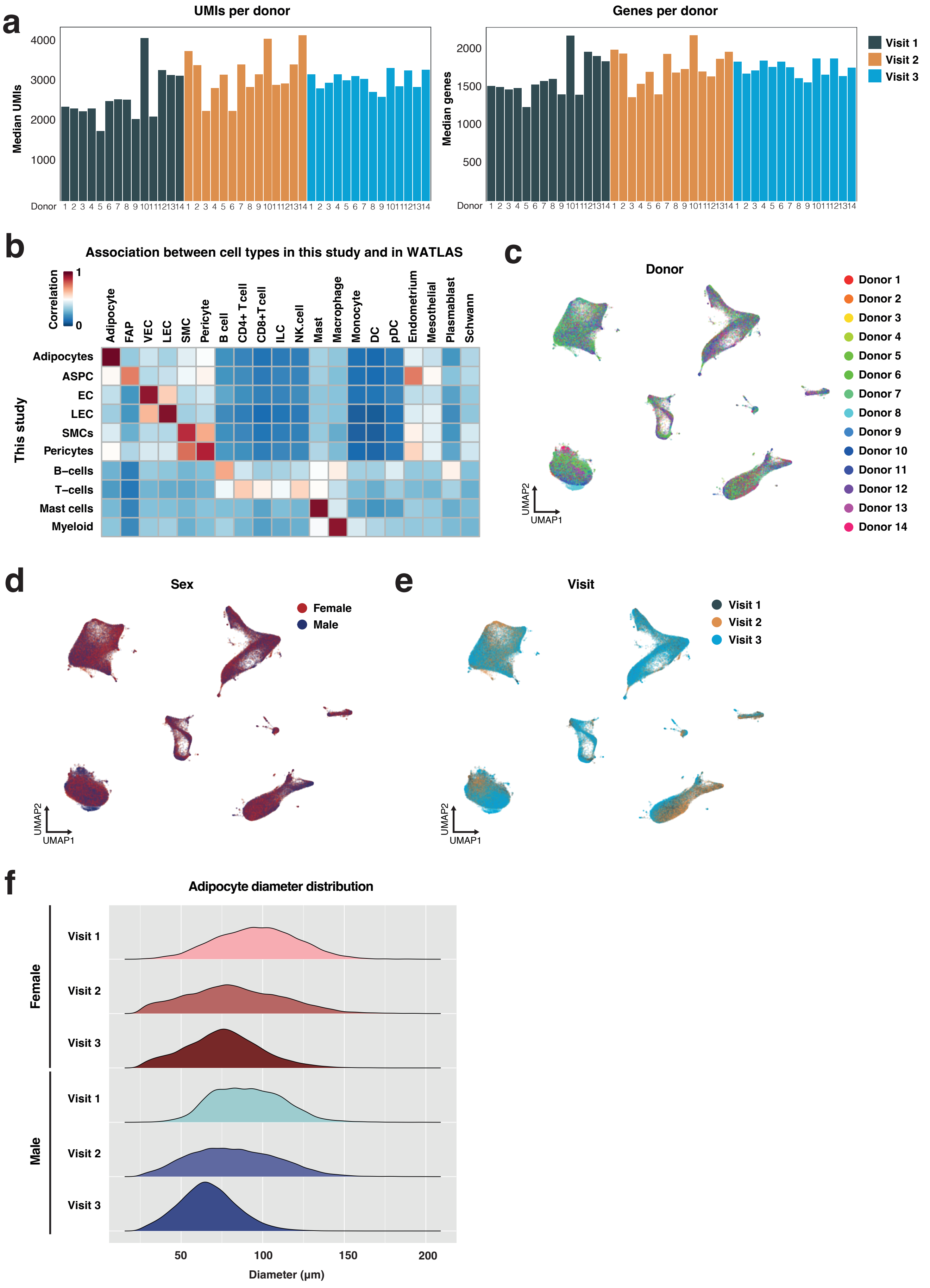

***Extended Data Fig. 1***

a, Median unique molecular identifiers (UMIs) (left) and median genes (right) detected for all 14 donors at each visit. b-d, Uniform manifold approximation and projection (UMAP) of 130,918 ASAT nuclei from all donors at all 3 visits with indication of the individual donors (b), sexes (c), and visits (d). e, Correlation of marker genes for cell types identified in this study with marker genes for cell types annotated in WATLAS [8]. f, The distribution of adipocyte diameters across visits, shown separately for each sex, is visualized using density ridges. This aggregated plot represents data summarized from 6 to 7 patients per condition.

### Extended Data Fig. 2

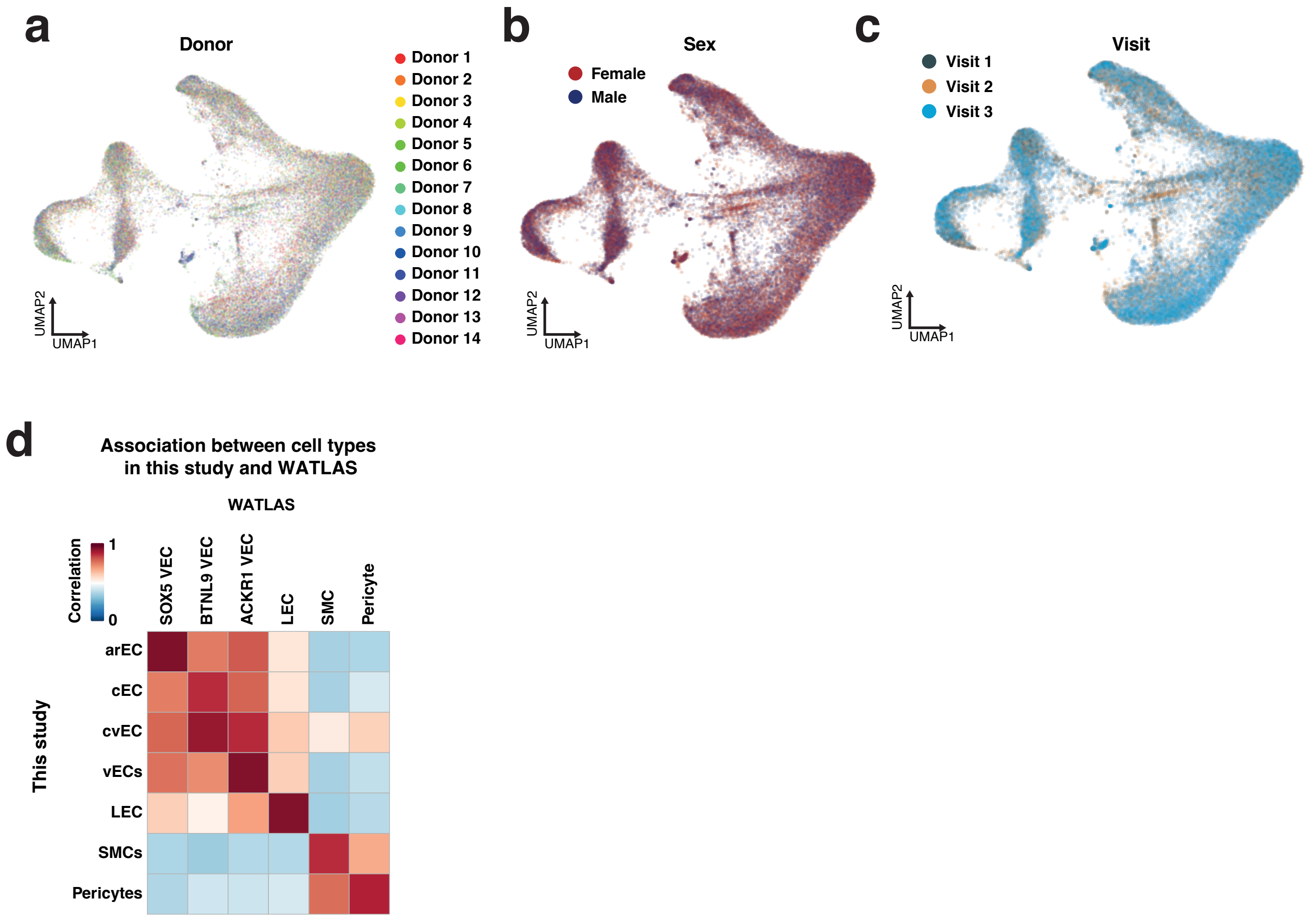

***Extended Data Fig. 2***

a-c, UMAP of vascular nuclei from all donors at all 3 visits with indication of the individual donors (a), sexes (b), and visits (c). d, Correlation of marker genes for vascular subpopulations identified in this study with marker genes for vascular subpopulations annotated in WATLAS [8].

### Extended Data Fig. 3

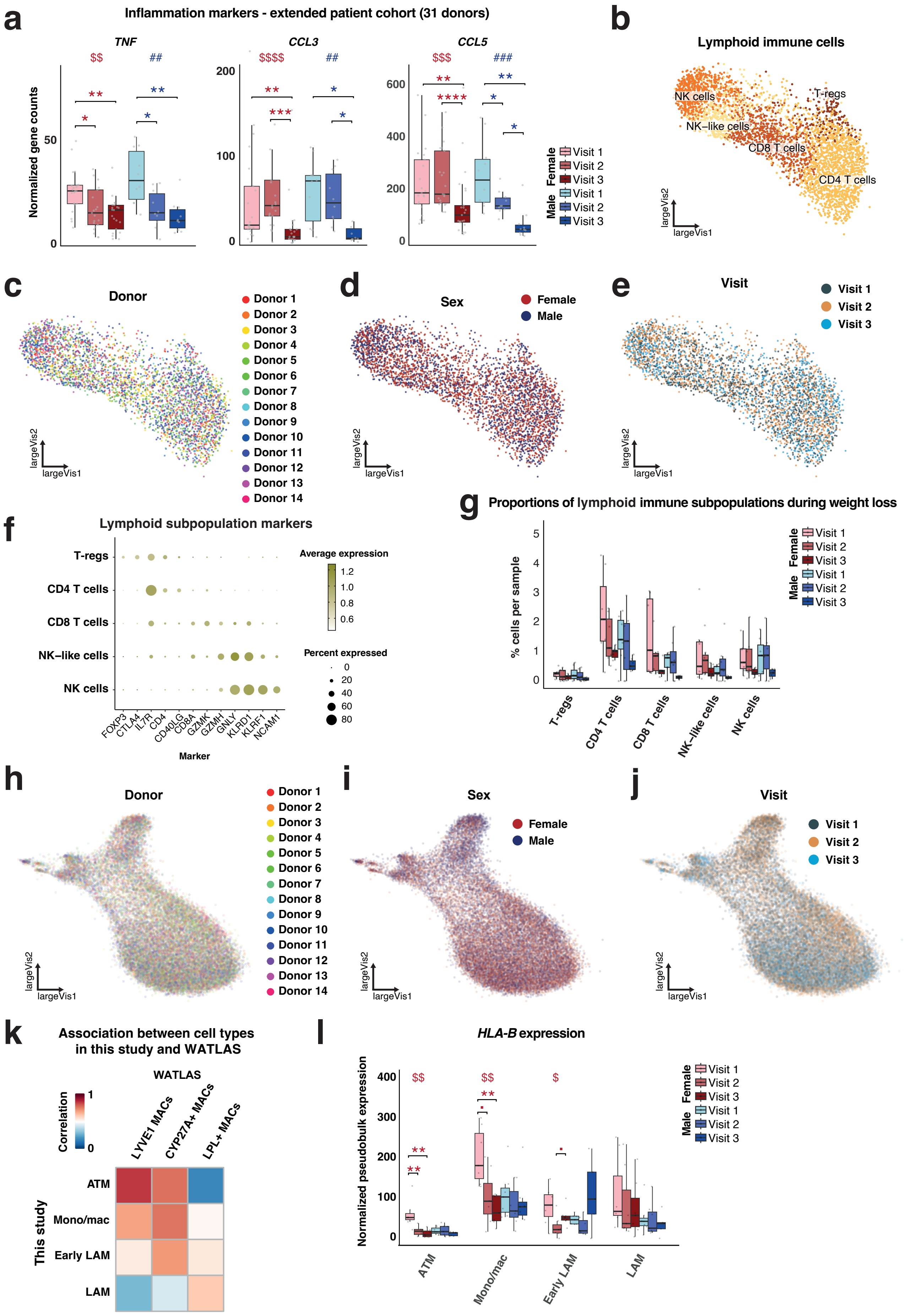

***Extended Data Fig. 3***

a, Normalized gene counts of indicated marker genes from the bulk RNA-seq data of 31 patients (female, n=21; male, n=10) at the indicated visits (due to y-axis constraints, a few data points for *TNF*, and *CCL3* were omitted from the graph). b-e, UMAP of lymphoid immune nuclei from all donors at all 3 visits with indication of subpopulations (b), the individual donors (c), sexes (d), and visits (e). f, Marker genes for each lymphoid immune subpopulation. g, The proportion of a specific lymphoid immune subpopulation relative to all cells in the human ASAT dataset. h-j, UMAP of myeloid immune nuclei from all donors at all 3 visits with indication of the individual donors (h), sexes (i), and visits (j). k, Correlation of marker genes for macrophage subpopulations identified in this study with marker genes for macrophage subpopulations annotated in WATLAS [8]. l, Normalized pseudobulk expression of *HLA-B* from the snRNA-seq data. Box plot data: center line, median; box limits, upper and lower quartiles; whiskers, 1.5x IQR. A KW test, followed by post-hoc Wilcoxon signed-rank tests (in a and g) or Wilcoxon rank-sum test (in l) with Holm's correction for multiple testing were performed to assess significance between visits for each sex separately (■, p<0.1; $/*, p<0.05; ##/$$/**, p<0.01; ###/$$$/***, p<0.001; $$$$/****, p<0.001). ATM, adipose tissue-resident macrophages; eLAMs, (early) lipid-associated macrophages ((e)LAMs); NK, naturel killer.

### Extended Data Fig. 4

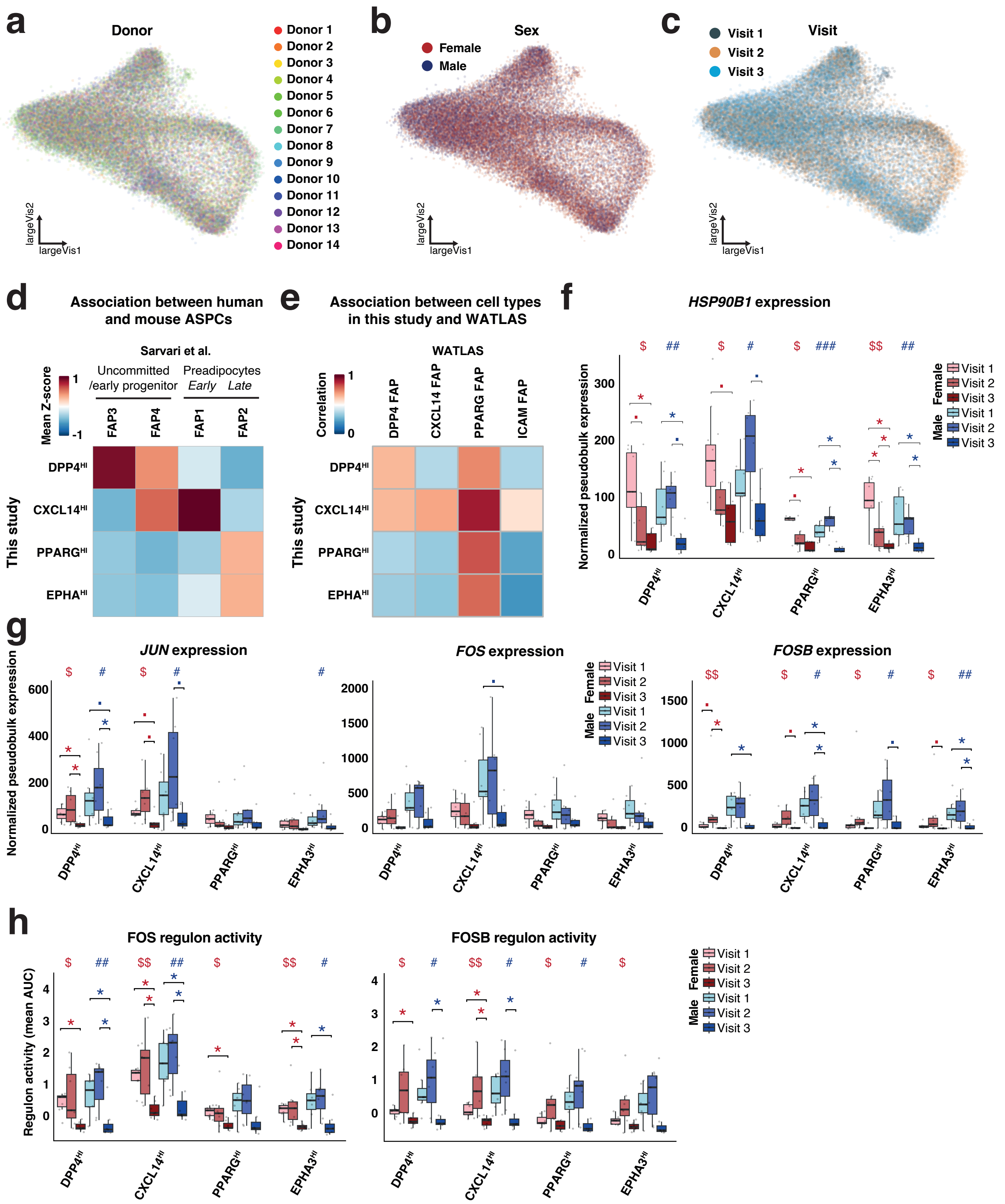

***Extended Data Fig. 4***

a-c, UMAP of ASPC nuclei from all donors at all 3 visits with indication of the individual donors (a), sexes (b), and visits (c). d-e, Correlation of marker genes for ASPCs identified in this study with marker genes for fibro-adipogenic progenitors annotated in Sarvari et al., [4] (d) or in WATLAS [8] (e). f, Normalized pseudobulk expression of *HSP90B1* from the snRNA-seq data. g, Normalized pseudobulk expression of AP1 transcription factors from the snRNA-seq data. h, Regulon activity (mean AUC) of FOS and FOSB. Box plot data: center line, median; box limits, upper and lower quartiles; whiskers, 1.5x IQR. A KW test, followed by a post-hoc Wilcoxon signed-rank test with Holm's correction for multiple testing, was performed to assess significance between visits for each sex separately in f, g, and h (■, p<0.1; #/$/*, p<0.05; ##/$$/**, p<0.01; $$$, p<0.001).

### Extended Data Fig. 5

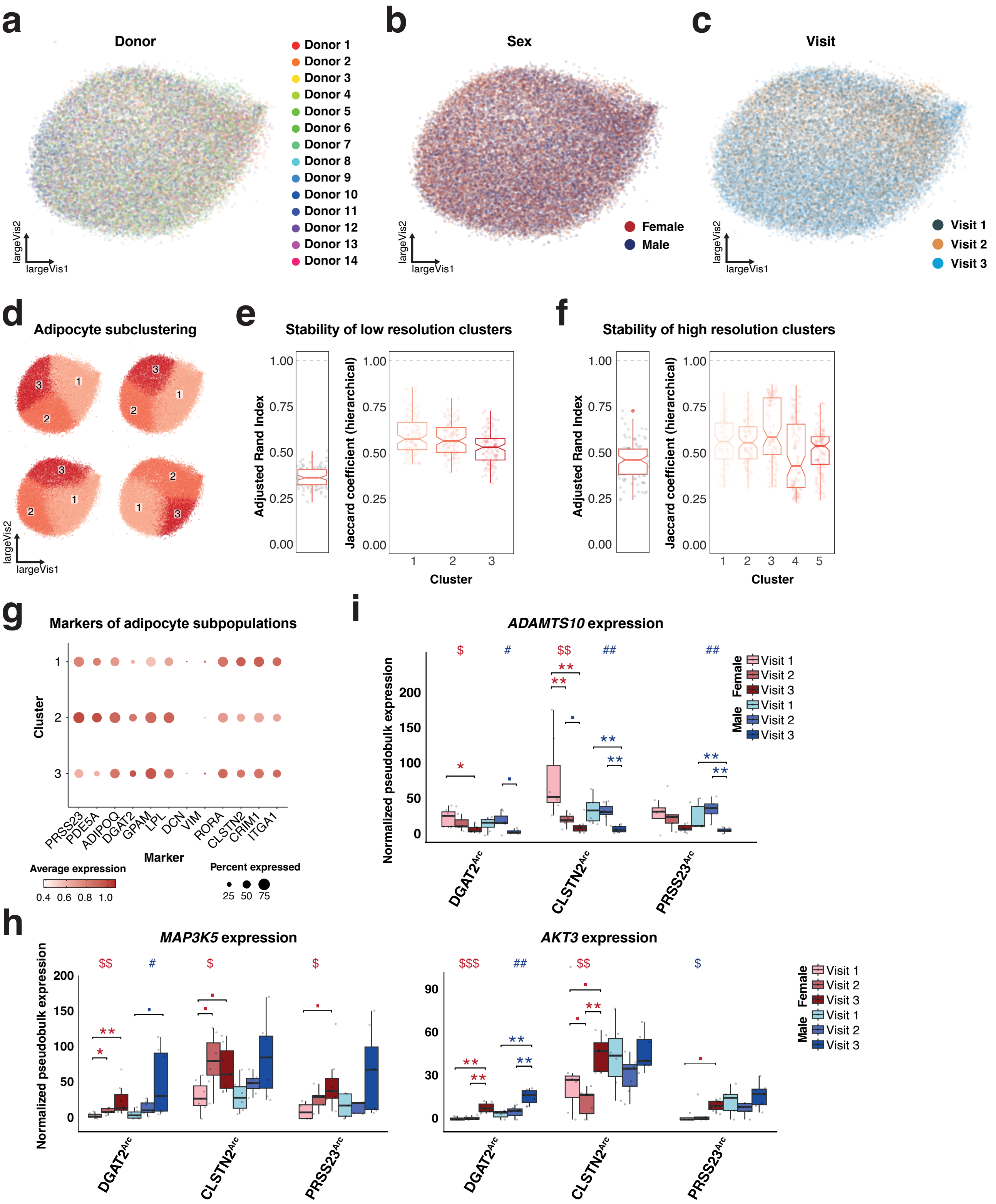

***Extended Data Fig. 5***

a-c, Uniform manifold approximation and projection (UMAP) of adipocyte nuclei from all donors at all 3 visits with indication of the individual donors (a), sexes (b), and visits (c). d, Repeated clustering analyses of adipocyte subpopulations using low resolution clustering. e-f, Cluster stability of adipocyte subpopulation using low resolution (e) or high resolution (f) clustering. g, Marker genes for the adipocyte subpopulation using low resolution clustering. h-i, Normalized pseudobulk expression of indicated cluster 3 (h) and cluster 4 (i) genes from the snRNA-seq data. Box plot data: center line, median; box limits, upper and lower quartiles; whiskers, 1.5x IQR. A KW test, followed by a post-hoc Wilcoxon signed-rank test with Holm's correction for multiple testing, was performed to assess significance between visits for each sex separately in h, and i (■, p<0.1; #/$/*, p<0.05; ##/$$/**, p<0.01).

### Extended Data Fig. 6

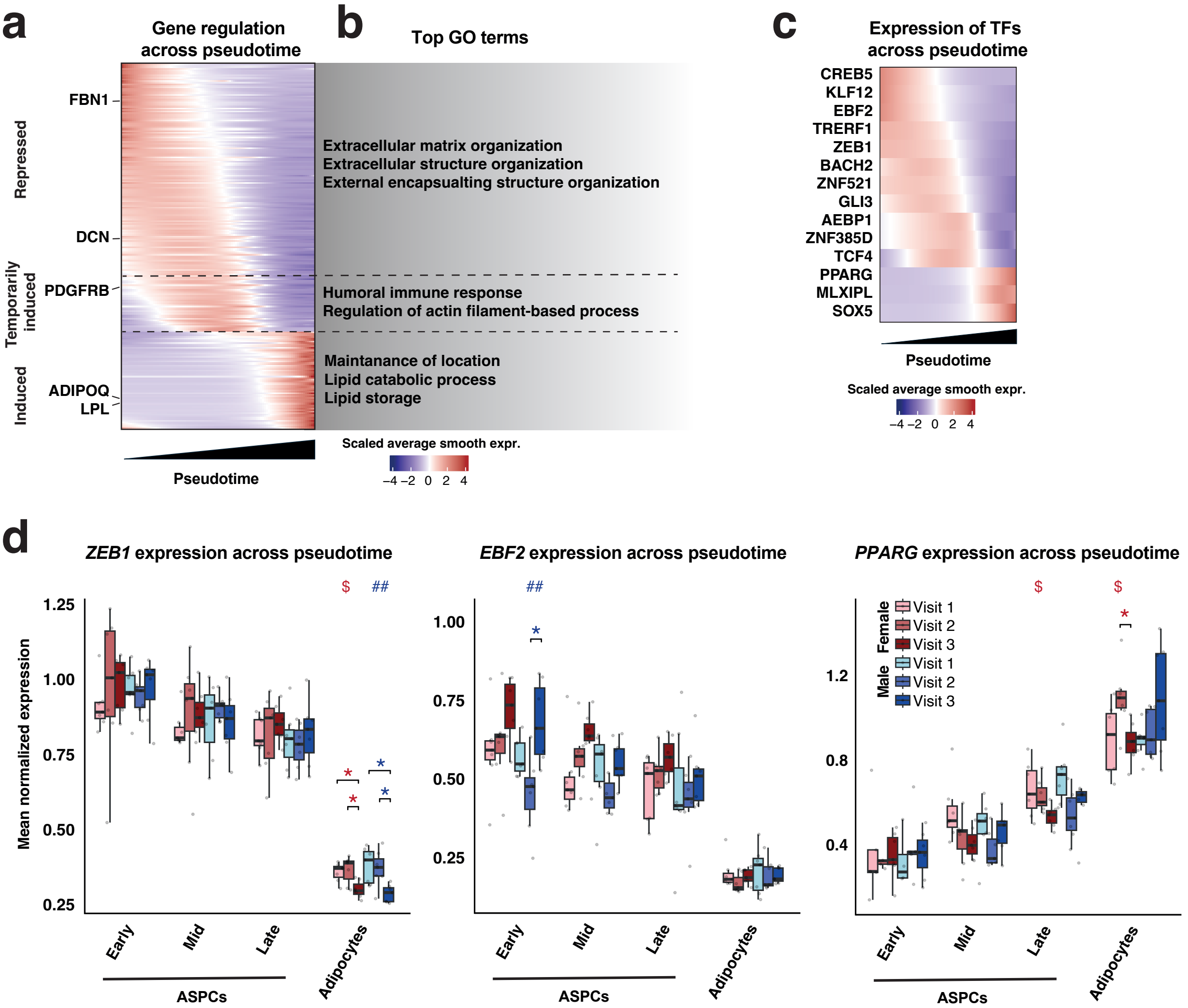

***Extended Data Fig. 6***

a, Heatmap showing average scaled expressions of genes regulated along pseudo-time across the adipogenic trajectory (in Fig. 6a-b) for all visits. b, the top GO term in each gene cluster (in a). c, Heatmap showing average scaled expressions of transcriptional regulators differentially expressed along pseudo-time across the trajectory (in Fig. 6a-b) for all visits. e) Expression of indicated genes encoding transcription factors at different cellular stages during adipocyte differentiation (as defined in 6b). Box plot data: center line, median; box limits, upper and lower quartiles; whiskers, 1.5x IQR. A KW test, followed by a post-hoc Wilcoxon rank-sum test with Holm's correction for multiple testing, was performed to assess significance between visits for each sex separately in d and ($/*, $p<0.05$; ##, $p<0.01$).

### Extended Data Fig. 7

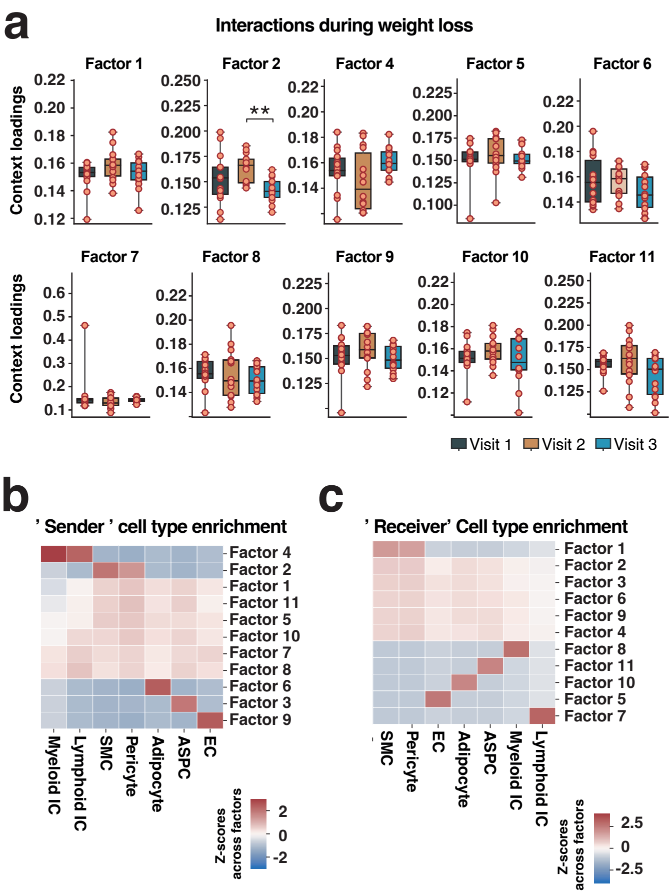

***Extended Data Fig. 7***

a, Sample loadings for each visit for each indicated factor as determined by LIANA and Tensor-cell2cell. b-c, Heatmaps displaying context loadings, represented as z-scores across all factors, for each cell type, highlighting their potential as either sender (b) or receiver (c) cell types in each factor. Box plot data: center line, median; box limits, upper and lower quartiles; whiskers, 1.5x IQR. A KW test, followed by a post-hoc Wilcoxon signed-ranked test with Holm's correction for multiple testing, was performed to assess significance between visits for each sex separately in a **, $p<0.01$.

**Extended data**

***Extended Data Table 1:*** Marker genes of all major cell types and subpopulations

***Extended Data Table 2:*** GO terms associated with marker genes of major cell types and subpopulations

***Extended Data Table 3:*** Macrophage DEG genes with indication of clusters and associated GO terms for each cluster.

***Extended Data Table 4:*** ASPC DEG genes with indication of clusters and associated GO terms for each cluster.

***Extended Data Table 5:*** Adipocyte archetype DEG genes with indication of clusters and associated GO terms for each cluster.

***Extended Data Table 6:*** Genes differentially regulated across the adipocyte differentiation trajectory

***Extended Video 1:*** Video displaying staining of F4/80 (red) from visit 1 donor.

***Extended Video 2:*** Video displaying staining of F4/80 (red), PLIN (green), and DAPI (blue) from visit 1 donor.
